# Body mass index and mortality in UK Biobank: revised estimates using Mendelian randomization

**DOI:** 10.1101/281436

**Authors:** Kaitlin H Wade, David Carslake, Naveed Sattar, George Davey Smith, Nicholas J Timpson

**Author notes:** **Corresponding author:** Dr Kaitlin H Wade, Oakfield House, Oakfield Road, Clifton, Bristol, BS8 2BN. **Disclosures**: NS is a member of the UK Biobank international scientific advisory board and enhancements committee. This had no bearing on the study. Otherwise, the authors declared no conflicts of interest. **Author Contributions**: All authors conceived and designed the study, interpreted results and approved the final manuscript. KHW had full access to all the data and performed all analyses with DC and takes responsibility for the integrity of the data and accuracy of the analysis. KHW, DC and NJT drafted the initial manuscript, the final version of which was critically appraised, revised and approved by all authors.

## Abstract

**Objective:** Obtain estimates of the causal relationship between different levels of body mass index (BMI) and mortality.

**Methods:** Mendelian randomization (MR) was conducted using genotypic variation reliably associated with BMI to test the causal effect of increasing BMI on all-cause and cause-specific mortality in participants of White British ancestry in UK Biobank.

**Results:** MR analyses supported existing evidence for a causal association between higher levels of BMI and greater risk of all-cause mortality (hazard ratio (HR) per 1kg/m^2^: 1.02; 95% CI: 0.97,1.06) and mortality from cardiovascular diseases (HR: 1.12; 95% CI: 1.02, 1.23), specifically coronary heart disease (HR: 1.19; 95% CI: 1.05, 1.35) and those other than stroke/aortic aneurysm (HR: 1.13; 95% CI: 0.93, 1.38), stomach cancer (HR: 1.30; 95% CI: 0.91, 1.86) and oesophageal cancer (HR: 1.08; 95% CI: 0.84, 1.38), and with decreased risk of lung cancer mortality (HR: 0.97; 95% CI: 0.84, 1.11). Sex-stratified analyses supported a causal role of higher BMI in increasing the risk of mortality from bladder cancer in males and other causes in females, but in decreasing the risk of respiratory disease mortality in males. The characteristic J-shaped observational association between BMI and mortality was visible with MR analyses but with a smaller value of BMI at which mortality risk was lowest and apparently flatter over a larger range of BMI.

**Conclusion:** Results support a causal role of higher BMI in increasing the risk of all-cause mortality and mortality from other causes. However, studies with greater numbers of deaths are needed to confirm the current findings.

## INTRODUCTION

Whilst severe obesity (a body mass index [BMI] ≥35kg/m^2^) clearly increases the risk of death, having a BMI above 25kg/m^2^ has been shown to increase the risk of all-cause mortality and mortality specifically from vascular diseases, diabetes, respiratory diseases and neoplastic (cancer) in a dose-response manner^1–7^. For example, each 5kg/m^2^ increase in BMI (or a transition between BMI categories for normal weight, overweight and obese, for example) has been shown to be associated with an increased risk of overall mortality by >30%, vascular mortality by 40% and the risk of diabetic, renal and hepatic mortality by 60-120%^1, 8^. It was also estimated that 3.6% (N∼481,000) of all new adult cancer cases in 2012 (aged >30 after a 10-year lag period) were estimated to be attributable to high BMI, a quarter of which could be attributed to the rise in BMI since 1982^9^. Furthermore, having excess weight from early life over the lifecourse increases the risk of later mortality^10–12^.

However, there have been inconsistencies within the current literature relating to the “obesity paradox”, whereby being overweight can appear seemingly protective^13–16^. Most prominently, in a systematic review and meta-analysis (providing a combined sample size of more than 2.88 million individuals), Flegal *et al*. showed a ∼6% lower risk of all-cause mortality in overweight individuals (i.e., a BMI of 25-30kg/m^2^) compared to normal weight individuals (i.e., a BMI of 18.5-25kg/m^2^)^13^. However, controversial findings like this are not without limitation, as confounding by age, ill-health and lifestyle factors plus selection bias (among other forms of bias) are likely^17–19^. Furthermore, many studies report a characteristic J-shaped curve in the association between BMI and the risk of mortality from varying causes^1, 2, 4, 6, 8, 16, 20–24^. In this context, individuals at the lower tail of the BMI distribution (i.e., underweight or below ∼22.5-25kg/m^2^) have an increased risk of mortality along with those above the ‘normal weight’ threshold^1, 2, 6, 8^. However, there are discrepancies in the reporting of this pattern, specifically between condition-specific mortality and populations of varying ancestries^4, 5, 25–27^.

Studies have used instrumental variable (IV) approaches to improve causal inference in the effect of exposures such as BMI, height and blood pressure (BP) on mortality using an intergenerational study design^28–30^. Using offspring exposures as IVs for the exposure of the parents in this context, whose mortality risk is being examined, provides a valid test as to whether an exposure (for example, BMI) has a causal effect on an outcome (for example, all-cause mortality) and allows the estimation of the magnitude of such an effect. Whilst this method has proved useful, there are caveats including, intergenerational confounding between the instrument (offspring exposure) and outcome (parental mortality) may lead to biased causal estimates.

Mendelian randomization (MR) is a well-documented extension of IV methodology that uses genetic variants (most commonly, single nucleotide polymorphisms [SNPs]) as IVs to provide a relatively unbiased causal estimate of the effect of an exposure (here, BMI) on an outcome (here, mortality)^31–34^. The use of MR has provided evidence to support a causal effect of higher BMI increasing the risk of coronary heart disease (CHD), stroke, aortic aneurysm and type-2 diabetes^35–39^; cardiometabolic traits such as BP, fasting glucose and insulin, C-reactive protein, interleukin-6 and cholesterol levels^40–42^ and various types of cancer including oesophageal, ovarian, lung, stomach, kidney, endometrial and colorectal cancer^43–52^, whilst decreasing the risk of Parkinson disease^53^ and breast cancer^43, 54^. However, there are inconsistencies within the field (for example, evidence suggesting no causal role of higher BMI in stroke^55^ or cancer risk^56^) and no study has explicitly used this technique to explore the causal role of BMI in all-cause and cause-specific mortality. Here, data from the UK Biobank study, a powerful and large resource of comprehensive phenotypic, genetic and death registry data from the UK, were used to generate estimates of causal role of BMI in both all-cause and cause-specific mortality, avoiding many of the problems of confounding and bias seen elsewhere.

## METHODS

### The UK Biobank study

The UK Biobank study recruited over 500,000 people aged 37-73 years (99.5% were between 40 and 69 years) from across the country in 2006-2010. Particularly focused on identifying determinants of human diseases in middle-aged and older individuals, participants provided a range of information (such as demographics, health status, lifestyle measures, cognitive testing, personality, self-report and physical/mental health measures) via questionnaires and interviews; anthropometric measures, BP readings and samples of blood, urine and saliva were taken. A full description of the study design, participants and quality control (QC) methods has been described in detail previously^57–59^. UK Biobank received ethical approval from the Research Ethics Committee (REC reference: 11/NW/0382).

Details of patient and public involvement in the UK Biobank are available online (www.ukbiobank.ac.uk/about-biobank-uk/ and https://www.ukbiobank.ac.uk/wp-content/uploads/2011/07/Summary-EGF-consultation.pdf?phpMyAdmin=trmKQlYdjj-nQIgj%2C-fAzikMhEnx6) and is available in a pre-print version^58^. No patients were specifically involved in setting the research question or the outcome measures, nor were they involved in developing plans for recruitment, design or implementation of this study. No patients were asked to advise on the interpretation or writing up of the results. There are no specific plans to disseminate the results of the research to study participants, but the UK Biobank disseminates key findings from the projects on its websites. At the time of this study, phenotypic data were available for 502,619 participants.

### Measures of body mass index

Weight and height were collected at baseline when participants attended the initial assessment centre. Height (cm) was measured using a Seca 202 device in all participants in the UK Biobank along with sitting height. Weight (kg) was measured by a variety of means during the initial Assessment Centre visit, which was amalgamated into a single weight variable on the UK Biobank release data.

A total of 13 participants had a height measurement more than 4.56 standard deviations (SDs) away from the mean and one person had a sitting to standing height ratio of greater than 0.75, which is not compatible with normal growth and development^60^. These participants were excluded, leaving 500,066 valid height measurements. Of these, 499,504 participants had weight measurements available (no weight values were excluded).

The UK Biobank currently has two different measures of adiposity – BMI calculated as weight divided by height squared (kg/m^2^) measured at the initial Assessment Centre visit and mass quantified using electrical impedance (in increments of 0.1kg), which was used to calculate a second measure of BMI. If BMI measured at the initial Assessment Centre visit was not available, the electrical impedance measure was used (n=255). Participants with substantial differences (>4.56SD) between impedance and normal BMI measures were excluded (n=1,164), if both measures were available. After these preliminary steps, 498,595 participants had a valid BMI measurement (see Figure 1 for flow-chart of the participants used in this analysis).

**Figure 1.**
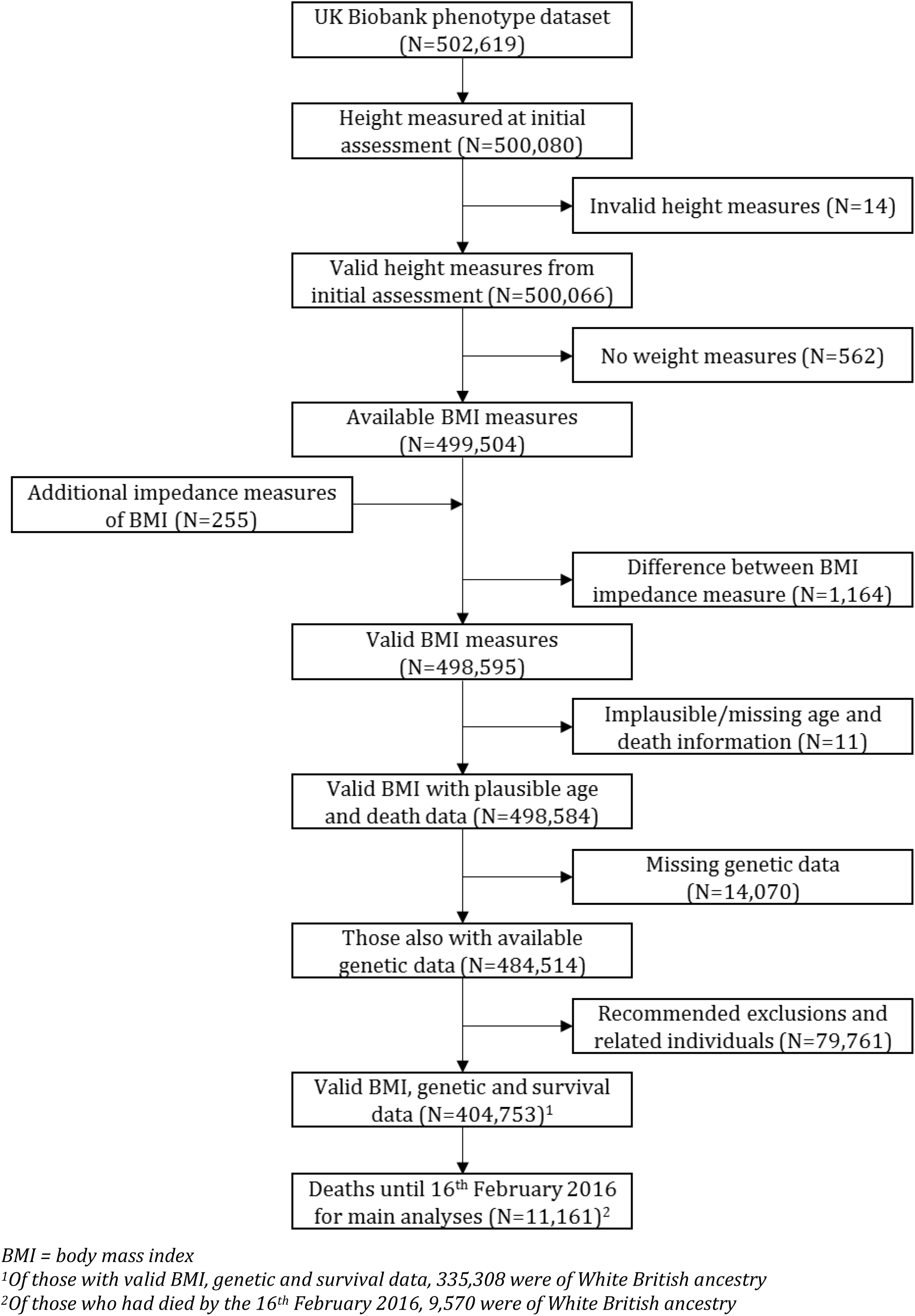
Flow-chart of those included in main analyses

### All-cause and cause-specific mortality

Data from death certificates were sent to UK Biobank on a quarterly basis provided by the National Health Service (NHS) Information Centre for participants from England and Wales and by NHS Central Register, Scotland for participants from Scotland. More detailed information on mortality are available at http://biobank.ctsu.ox.ac.uk/crystal/refer.cgi?id=115559. The death certificates include the disease or condition stated to be the underlying cause of death, as well as other conditions, diseases, injuries or events contributing to death but not related to the disease or condition causing it. Data were provided as date of death (DoD), an integer value for age of death (AoD) and underlying (primary) cause of death in International Classification of Diseases (ICD)-10 codes for all deaths that occurred between the 10/05/2006 and 16/02/2016. Rather than using the integer value of AoD from the death certificate, a more precise measure of AoD was derived by adding the time interval between date of initial assessment and DoD (in days) to the participant’s age at initial assessment. All participants who were not recorded as dead by the 16^th^ of February 2016 were assumed to still be alive. The ICD-10 codes were categorised into all-cause and cause-specific mortality as presented in Table S1a. As of August 2017 (date of extraction for all data), there were 14,417 total deaths in the entire UK Biobank dataset (Table S1a), which remains to be the most updated data on mortality.

### Covariables

At the initial UK Biobank Assessment Centre, participants were given a touchscreen questionnaire, which included questions about sociodemographic status, early life, sex-specific factors, lifestyle and environment, family history, health and medical history and psychosocial factors. Of the sociodemographic questions, participants were asked whether they had any of the following qualifications or equivalent: i) college or university degree, ii) A/AS-levels, iii) O-levels/GCSEs, iv) CSEs, v) NVQ or HND or HNC, vi) other professional qualifications eg. nursing or teaching, vii) none of the listed. Additionally, participants were asked which of the following described their current employment situation: i) in paid employment or self-employed, ii) retired, iii) looking after family home and/or family, iv) unable to work because of sickness or disability, v) unemployed, vi) doing unpaid or voluntary work, vii) full or part-time student, viii) none of the listed. Answers to these questions were used to derive variables the represented the participants’ highest qualification level and current employment status, respectively.

Of the lifestyle and environment questions, participants were asked their smoking and alcohol drinking status, categorised into ‘never’, ‘former or ‘current’. Participants were also asked how many days in a typical week they would do 10 or more minutes of vigorous physical activity (“activities that make you sweat or breathe hard such as fast cycling, aerobic exercise and heavy lifting”).

### Genotyping

Pre-imputation, QC, phasing and imputation of UK Biobank have been described elsewhere^58, 61^. The genetic variants used were extracted genotypes from the UK Biobank imputation dataset (using only genetic variants imputed to the Haplotype Reference Consortium (HRC) reference panel), which had extensive QC performed including exclusion of the majority of third degree or closer relatives from a genetic kinship analysis of 96% of participants. For more details, see http://biobank.ctsu.ox.ac.uk. A total of 77 common genetic variants associated with BMI within people of only European ancestry (and excluding those that reached genome-wide levels of statistical confidence in only one sex or one stratum) in the most updated genome-wide association study (GWAS) conducted by the GIANT consortium (comprising up to 339,224 people) were extracted for MR analyses (Table S2)^60, 62^. One SNP from the GWAS (rs12016871) was not present in the UK Biobank imputed genetic data, so a proxy SNP (i.e., one that is in linkage disequilibrium [LD] with rs12016871) was identified (rs4771122; r^2^=0.876, distance=2398bp) and used in replacement^62^. We checked that each of the variants were imputed with high quality (>0.90, Table S2).

The dosage of each genetic variant was weighted by its relative effect size on BMI obtained from the GIANT consortium^62^ and summed across all variants. The resulting total was then rescaled by dividing by the sum of all effect sizes and multiplied by the number of genetic variants used. Therefore, this weighted genetic risk score (GRS) reflected the average number of BMI-increasing alleles each participant possessed^60^. In total, 487,409 participants had genetic data.

### Statistical analysis

As only the month and year of birth was available in the UK Biobank study, date of birth (DoB) was set as the 15^th^ of each month and year in which the participant was born. Participants were removed if they lacked information on date of birth (used for secular trends), initial assessment age and date, cause of death or AoD. Participants were also excluded if they lacked any/plausible information on DoD (i.e., if the individual had apparently died before the assessment clinic that they later attended). Participants who were never at risk during the follow-up period (i.e., who were recruited after 16^th^ February 2016) were also excluded.

The sample was restricted to UK Biobank participants of White British ancestry, defined by those who self-identified as “White British” and confirmed to have a similar genetic ancestry based on a principal components (PCs) analysis of the genome-wide genetic data^58^. Of those with full genetic data and information on BMI, 335,308 participants were included in analyses after recommended exclusions based on sex mismatch, sex-chromosome aneuploidy detection and related individuals (Supporting Methods, Figure 1). Of these, 9,570 had available data on cause, age and date of death (Table S1a, Figure 1).

Within the set of participants who had required information, Cox proportional hazards regression models were used to estimate conventional hazard ratios (HRs) for all-cause and cause-specific mortality per unit increase (kg/m^2^) of BMI. The participant’s age was used as a measure of time; thus, models were automatically adjusted for age. Analyses were conducted with the following two models: i) unadjusted and ii) adjusted for secular trends (DoB), current occupation, qualifications, smoking status, alcohol intake, physical activity and the first ten genetic PCs. Analyses were restricted to the conditions responsible a minimum number of deaths (>40)^29^ and performed in the whole sample plus stratified by sex. The associations of the GRS with BMI and of each covariable with BMI and the GRS were tested using linear regression and associations of each covariable with all-cause mortality was assessed using Cox proportional hazards regression models, with all analyses adjusting for the first ten genetic PCs.

For MR analyses, a two-stage approach was conducted within the context of survival analyses. In the first stage, BMI was regressed on a GRS comprising 77 SNPs to generate the denominator of a two-stage MR estimate. In the second stage, Cox proportional hazards models were used to estimate the HR of each mortality outcome with each unit increase in the GRS, thus generating the numerator for the two-stage MR estimate. Exponentiating the ratio between the natural logarithm of the numerator (HR for each mortality outcome per unit increase in the GRS) and the denominator (association between BMI and the GRS) yielded a ratio MR estimate of the HR of each mortality outcome per unit increase (kg/m^2^) in BMI. Confidence intervals (CIs) were obtained using Taylor series expansions^63^. A simplification of the matrix method for the Durbin-Wu-Hausman (DWH) test for endogeneity was used to compare the HR estimates per unit increase in BMI derived from conventional Cox regression and the two-stage MR estimate (see Supporting Methods). All analyses were conducted using Stata 14.2.

### Linearity and proportional hazards assumption

To test the linearity of associations, fully-adjusted cubic spline models for both the exposure (here, BMI) and the instrument (here, the GRS) were plotted to show the pattern of association between levels of BMI and mortality. Observations were censored to restrict them to the period when death would have been recorded and used in the analysis. To focus on the densest part of the BMI distribution, the linearity tests were conducted after removing data below/above the 1^st^/99^th^ percentile, respectively, due to the scarcity of data towards the tails of the BMI distribution. In addition, an approximate MR analogue to the non-linear plot of mortality against BMI was obtained by estimating localized average causal effects (i.e., MR estimates of the log-linear effect of BMI on the hazard of mortality) within percentiles of the instrument-free exposure (set at the 5^th^, 10^th^, 25^th^, 50^th^, 75^th^ and 85^th^ percentile)^64^. These localised average causal effects were then joined together and plotted against corresponding quantiles of the original exposure^65^. HRs were calculated relative to the mean BMI of 27kg/m^2^ and 1000 bootstrap resamples were used to obtain CIs. Meta-regression was used to test for a linear trend in association between the GRS and BMI (i.e. denominator of the two-stage MR estimate) over quantiles of the instrument-free exposure.

To check the proportional hazards assumption, the Schoenfeld residuals for BMI from the cubic spline models of each mortality outcome were tested for association with the natural log of the follow-up time (here, age) using both conventional Cox regression and MR analysis. If there was evidence for a difference in the association between BMI and the risk of mortality with age (i.e., a Bonferroni-corrected P-value of 0.05/138 tests = 0.0004), an interaction term was fitted to the cubic spline model using the “tvc()” option for survival analyses in Stata.

### Sensitivity analyses

To investigate the validity of the GRS as an IV within this context, MR-Egger was used to detect and accommodate violations of the MR assumptions, specifically horizontal pleiotropy^66^. The intercept obtained from the MR-Egger test is used as an indication of pleiotropy and the slope can be considered as the estimate of the causal effect between the exposure (here, BMI) and the outcome (here, all-cause and cause-specific mortality). In addition, the weighted median- and mode-based methods were used^67, 68^, which vary in their assumptions of instrument validity. The weighted median approach provides a causal estimate even when 50% of instruments are invalid and the weighted mode estimate is consistent when the largest number of similar causal effect estimates comes from valid instruments, even if most instruments are invalid. MR-Egger, weighted median and weighted mode estimates were compared to those obtained from the inverse-variance weighted (IVW) for two-sample MR^66^. For these analyses, the first-stage estimates (coefficients of the association between each SNP and BMI) were obtained from an independent external source, as to not induce weak instrument bias^69, 70^, and the second-stage estimates (natural logarithm of the HR for each mortality outcome with each SNP) were obtained directly from UK Biobank.

In the UK Biobank sample, there is evidence to suggest a differential array effect on markers scattered across the genome (i.e., those that were genotyped using the Affymetrix UK Biobank Axiom® Array or the Affymetrix UK BiLEVE Axiom Array^58, 61^) and the UK BiLEVE sub-sample, which included >50,000 participants and used the UK BiLEVE Axiom Array, also preferentially selected individuals based on smoking intensity. To evaluate the impact of this differential array effect, MR sensitivity analyses were conducted with an additional adjustment of genotyping chip.

As a final sensitivity analysis, the GRS was restricted to exclude the genetic variants known to be classified as having a secondary signal within a locus to other phenotypes (N=7; leaving 70 in the GRS, Table S3)^37, 60^ and all MR analyses were repeated.

## RESULTS

Of those included, the average age of initial assessment was 56.9 years old (SD=8) and participants had an average BMI of 27.4kg/m^2^ (SD=4.7) (Table 1). Of the 335,308 participants with required information for mortality analyses, 9,570 participants (N=5,882/3,688 males/females, respectively) had died by the 16^th^ of February 2016 at an average age of 65.7 years old (SD=6.9). Most had died from various types of cardiovascular diseases (CVDs) and cancer (Table S1a/1b for whole-sample and sex-specific mortality).

**Table 1.** Descriptive statistics for UK Biobank participants of White British Ancestry included in the main analyses

| **Variable** | **N** | **Mean (SD) or percentage** |
| --- | --- | --- |
| Age (years) at initial assessment | 335,308 | 56.87 (8.00) |
| Sex (% of males) | 335,308 | 46.22 |
| Body mass index (kg/m<sup>2</sup>) | 335,308 | 27.38 (4.74) |
| *Smoking status* | 334,142 |  |
| Never | 183,170 | 54.82 |
| Former | 117,838 | 35.26 |
| Current | 33,134 | 9.92 |
| *Alcohol drinker status* | 335,074 |  |
| Never | 10,311 | 3.08 |
| Former | 11,368 | 3.39 |
| Current | 313,395 | 93.53 |
| *Highest qualifications* | 275,544 |  |
| College or University degree | 106,280 | 38.57 |
| A-levels | 38,271 | 13.89 |
| O-levels | 73,770 | 26.77 |
| CSEs | 18,016 | 6.54 |
| NVQ/HND/HNC | 22,012 | 7.99 |
| Other professional qualifications | 17,195 | 6.24 |
| *Current employment status* | 332,835 |  |
| In paid employment or self-employed | 190,085 | 57.11 |
| Retired | 117,615 | 35.34 |
| Looking after home/family | 8,690 | 2.61 |
| Unable to work due to sickness/disability | 9,982 | 3.00 |
| Unemployed | 4,436 | 1.33 |
| Doing unpaid or voluntary work | 1,404 | 0.42 |
| Full or part-time student | 623 | 0.19 |
| Days/week spent doing vigorous physical activity | 319,813 | 1.82 (1.94) |
| Genotyping chip<sup>2</sup> | 335,308 | 9.24 |
| Age at death (years) | 9,570 | 65.66 (6.88) |
| Date of death<sup>1</sup> |  | 06/02/2013<br>(07/07/2007-16/02/2016) |

CSE = certificate of secondary education; HNC = higher national certificate; HND = higher national diploma; NVQ = national vocational qualification; SD = standard deviation
<sup>1</sup>Recorded as the mean (minimum and maximum) date of death
<sup>2</sup>UK BiLEVE participants genotyped on the Affymetrix Axiom Array

### Observational analyses

In unadjusted observational analyses, higher BMI was associated with a higher risk of all-cause mortality and mortality from CVD, specifically CHD and those other than stroke and aortic aneurysm, overall cancer and cancer in the stomach, oesophagus, kidney and liver (Table 2). Higher BMI was also associated with a lower risk of lung cancer mortality.

**Table 2.**
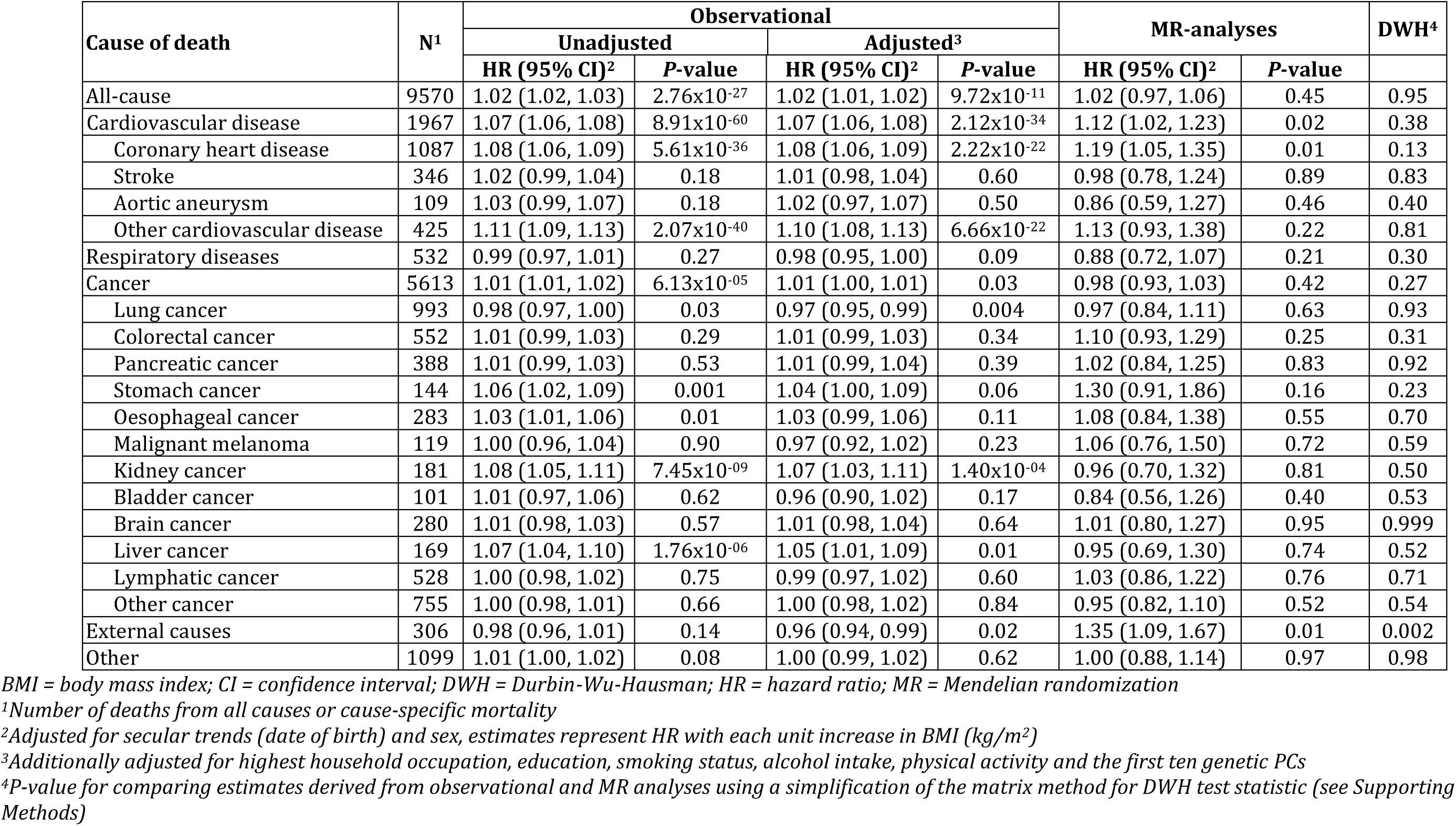
Observational and MR analyses of all-cause and cause-specific mortality by BMI in UK Biobank participants of White British ancestry

| Cause of death | N<sup>1</sup> | Observational |  |  |  | MR-analyses |  | DWH<sup>4</sup> |
| --- | --- | --- | --- | --- | --- | --- | --- | --- |
|  |  | Unadjusted |  | Adjusted<sup>3</sup> |  |  |  |  |
|  |  | HR (95% CI)<sup>2</sup> | P-value | HR (95% CI)<sup>2</sup> | P-value | HR (95% CI)<sup>2</sup> | P-value |  |
| All-cause | 9570 | 1.02 (1.02, 1.03) | 2.76x10<sup>-27</sup> | 1.02 (1.01, 1.02) | 9.72x10<sup>-11</sup> | 1.02 (0.97, 1.06) | 0.45 | 0.95 |
| Cardiovascular disease | 1967 | 1.07 (1.06, 1.08) | 8.91x10<sup>-60</sup> | 1.07 (1.06, 1.08) | 2.12x10<sup>-34</sup> | 1.12 (1.02, 1.23) | 0.02 | 0.38 |
| Coronary heart disease | 1087 | 1.08 (1.06, 1.09) | 5.61x10<sup>-36</sup> | 1.08 (1.06, 1.09) | 2.22x10<sup>-22</sup> | 1.19 (1.05, 1.35) | 0.01 | 0.13 |
| Stroke | 346 | 1.02 (0.99, 1.04) | 0.18 | 1.01 (0.98, 1.04) | 0.60 | 0.98 (0.78, 1.24) | 0.89 | 0.83 |
| Aortic aneurysm | 109 | 1.03 (0.99, 1.07) | 0.18 | 1.02 (0.97, 1.07) | 0.50 | 0.86 (0.59, 1.27) | 0.46 | 0.40 |
| Other cardiovascular disease | 425 | 1.11 (1.09, 1.13) | 2.07x10<sup>-40</sup> | 1.10 (1.08, 1.13) | 6.66x10<sup>-22</sup> | 1.13 (0.93, 1.38) | 0.22 | 0.81 |
| Respiratory diseases | 532 | 0.99 (0.97, 1.01) | 0.27 | 0.98 (0.95, 1.00) | 0.09 | 0.88 (0.72, 1.07) | 0.21 | 0.30 |
| Cancer | 5613 | 1.01 (1.01, 1.02) | 6.13x10<sup>-05</sup> | 1.01 (1.00, 1.01) | 0.03 | 0.98 (0.93, 1.03) | 0.42 | 0.27 |
| Lung cancer | 993 | 0.98 (0.97, 1.00) | 0.03 | 0.97 (0.95, 0.99) | 0.004 | 0.97 (0.84, 1.11) | 0.63 | 0.93 |
| Colorectal cancer | 552 | 1.01 (0.99, 1.03) | 0.29 | 1.01 (0.99, 1.03) | 0.34 | 1.10 (0.93, 1.29) | 0.25 | 0.31 |
| Pancreatic cancer | 388 | 1.01 (0.99, 1.03) | 0.53 | 1.01 (0.99, 1.04) | 0.39 | 1.02 (0.84, 1.25) | 0.83 | 0.92 |
| Stomach cancer | 144 | 1.06 (1.02, 1.09) | 0.001 | 1.04 (1.00, 1.09) | 0.06 | 1.30 (0.91, 1.86) | 0.16 | 0.23 |
| Oesophageal cancer | 283 | 1.03 (1.01, 1.06) | 0.01 | 1.03 (0.99, 1.06) | 0.11 | 1.08 (0.84, 1.38) | 0.55 | 0.70 |
| Malignant melanoma | 119 | 1.00 (0.96, 1.04) | 0.90 | 0.97 (0.92, 1.02) | 0.23 | 1.06 (0.76, 1.50) | 0.72 | 0.59 |
| Kidney cancer | 181 | 1.08 (1.05, 1.11) | 7.45x10<sup>-09</sup> | 1.07 (1.03, 1.11) | 1.40x10<sup>-04</sup> | 0.96 (0.70, 1.32) | 0.81 | 0.50 |
| Bladder cancer | 101 | 1.01 (0.97, 1.06) | 0.62 | 0.96 (0.90, 1.02) | 0.17 | 0.84 (0.56, 1.26) | 0.40 | 0.53 |
| Brain cancer | 280 | 1.01 (0.98, 1.03) | 0.57 | 1.01 (0.98, 1.04) | 0.64 | 1.01 (0.80, 1.27) | 0.95 | 0.999 |
| Liver cancer | 169 | 1.07 (1.04, 1.10) | 1.76x10<sup>-06</sup> | 1.05 (1.01, 1.09) | 0.01 | 0.95 (0.69, 1.30) | 0.74 | 0.52 |
| Lymphatic cancer | 528 | 1.00 (0.98, 1.02) | 0.75 | 0.99 (0.97, 1.02) | 0.60 | 1.03 (0.86, 1.22) | 0.76 | 0.71 |
| Other cancer | 755 | 1.00 (0.98, 1.01) | 0.66 | 1.00 (0.98, 1.02) | 0.84 | 0.95 (0.82, 1.10) | 0.52 | 0.54 |
| External causes | 306 | 0.98 (0.96, 1.01) | 0.14 | 0.96 (0.94, 0.99) | 0.02 | 1.35 (1.09, 1.67) | 0.01 | 0.002 |
| Other | 1099 | 1.01 (1.00, 1.02) | 0.08 | 1.00 (0.99, 1.02) | 0.62 | 1.00 (0.88, 1.14) | 0.97 | 0.98 |

BMI = body mass index; CI = confidence interval; DWH = Durbin-Wu-Hausman; HR = hazard ratio; MR = Mendelian randomization
<sup>1</sup>Number of deaths from all causes or cause-specific mortality
<sup>2</sup>Adjusted for secular trends (date of birth) and sex, estimates represent HR with each unit increase in BMI (kg/m<sup>2</sup>)
<sup>3</sup>Additionally adjusted for highest household occupation, education, smoking status, alcohol intake, physical activity and the first ten genetic PCs
<sup>4</sup>P-value for comparing estimates derived from observational and MR analyses using a simplification of the matrix method for DWH test statistic (see Supporting Methods)

Adjusting Cox regression models for covariables made little difference to results, providing evidence that BMI was associated with a higher risk of all-cause mortality (HR per 1kg/m^2^ higher BMI: 1.02; 95% CI: 1.01, 1.02; *P*=9.72×10^−11^) and mortality from CVD (HR: 1.07; 95% CI: 1.06, 1.08; *P*=2.12×10^−34^), specifically CHD (HR: 1.08; 95% CI: 1.06, 1.09; *P*=2.22×10^−22^) and those other than stroke and aortic aneurysm (HR: 1.10; 95% CI: 1.08, 1.13; *P*=6.66×10^−22^), as well as a higher risk of mortality from overall cancer (HR: 1.01; 95% CI: 1.00, 1.01; *P*=0.03) and cancer of the kidney (HR: 1.07; 95% CI: 1.03, 1.11; *P*=1.40×10^−04^) and liver (HR: 1.05; 95% CI: 1.01, 1.09; *P*=0.01) (Table 2). The inverse association between BMI and risk of lung cancer also remained consistent after adjusting for covariables (HR: 0.97; 95% CI: 0.95, 0.99; *P*=0.004) and there was evidence for an inverse association between BMI and mortality from external causes (HR: 0.96; 95% CI: 0.94, 0.99; *P*=0.02). The association between BMI and risk of mortality from stomach cancer slightly attenuated (HR: 1.04; 95% CI: 1.00, 1.09; *P*=0.06) and, whilst the estimate of association between BMI and risk of mortality from oesophageal cancer remained consistent (HR: 1.03; 95% CI: 0.99, 1.06; *P*=0.11), CIs widened after adjustment (Table 2).

Results were similar within sex-stratified analyses, with additional evidence for an association between higher BMI and decreased risk of mortality from respiratory disease in males (HR: 0.91; 95% CI: 0.88, 0.95; *P*=2.28×10^−06^) but an increased risk of mortality from respiratory disease in females (HR: 1.06; 95% CI: 1.02, 1.09; *P*=0.002), which was not present in the overall sample (Table 3 and Table 4 for males and females, respectively). In males, adjusted Cox regression analyses also provided evidence for an association between higher BMI and increased risk of prostate cancer mortality (HR: 1.05; 95% CI: 1.02, 1.08; *P*=0.003), alongside greater magnitudes of association of higher BMI with a decreased risk of mortality from lung cancer (HR: 0.94; 95% CI: 0.91, 0.97; *P*=3.54×10^−05^) and bladder cancer (HR: 0.93; 95% CI: 0.87, 1.00; *P*=0.06), and increased risk of mortality from oesophageal cancer (HR: 1.07; 95% CI: 1.04, 1.11; *P*=8.50×10^−05^) and liver cancer (HR: 1.08; 95% CI: 1.03, 1.13; *P*=0.003). The estimate of association between higher BMI and mortality from external causes was smaller compared to that in the whole sample (HR: 0.97; 95% CI: 0.93, 1.01; *P*=0.11); however, all CIs overlapped (Table 3).

**Table 3.** Observational and MR analyses of all-cause and cause-specific mortality by BMI in males of White British ancestry

| Cause of death | N<sup>1</sup> | Observational |  |  |  | MR-analyses |  | DWH<sup>4</sup> |
| --- | --- | --- | --- | --- | --- | --- | --- | --- |
|  |  | Unadjusted |  | Adjusted<sup>3</sup> |  | HR (95% CI)<sup>2</sup> | P-value |  |
|  |  | HR (95% CI)<sup>2</sup> | P-value | HR (95% CI)<sup>2</sup> | P-value |  |  |  |
| All-cause | 5882 | 1.03 (1.02, 1.03) | 4.00x10<sup>-18</sup> | 1.02 (1.01, 1.03) | 2.65x10<sup>-07</sup> | 1.03 (0.97, 1.09) | 0.39 | 0.84 |
| Cardiovascular disease | 1467 | 1.08 (1.07, 1.09) | 1.23x10<sup>-48</sup> | 1.08 (1.06, 1.09) | 1.12x10<sup>-27</sup> | 1.10 (0.98, 1.24) | 0.10 | 0.67 |
| Coronary heart disease | 906 | 1.08 (1.07, 1.09) | 8.84x10<sup>-32</sup> | 1.08 (1.06, 1.09) | 8.61x10<sup>-18</sup> | 1.20 (1.04, 1.40) | 0.02 | 0.14 |
| Stroke | 194 | 1.03 (0.99, 1.06) | 0.10 | 1.02 (0.98, 1.07) | 0.27 | 0.90 (0.64, 1.26) | 0.52 | 0.43 |
| Aortic aneurysm | 83 | 1.04 (0.99, 1.09) | 0.14 | 1.03 (0.96, 1.09) | 0.41 | 0.82 (0.52, 1.31) | 0.40 | 0.34 |
| Other cardiovascular disease | 284 | 1.11 (1.09, 1.14) | 9.96x10<sup>-25</sup> | 1.11 (1.08, 1.14) | 9.33x10<sup>-15</sup> | 1.06 (0.82, 1.37) | 0.66 | 0.72 |
| Respiratory diseases | 361 | 0.94 (0.91, 0.97) | 1.08x10<sup>-05</sup> | 0.91 (0.88, 0.95) | 2.28x10<sup>-06</sup> | 0.84 (0.65, 1.09) | 0.19 | 0.53 |
| Cancer | 3113 | 1.01 (1.00, 1.02) | 0.002 | 1.01 (1.00, 1.02) | 0.06 | 1.00 (0.92, 1.08) | 0.95 | 0.76 |
| Lung cancer | 571 | 0.96 (0.94, 0.98) | 2.28x10<sup>-04</sup> | 0.94 (0.91, 0.97) | 3.54x10<sup>-05</sup> | 0.96 (0.79, 1.18) | 0.70 | 0.82 |
| Prostate cancer | 308 | 1.03 (1.01, 1.06) | 0.01 | 1.05 (1.02, 1.08) | 0.003 | 0.73 (0.57, 0.93) | 0.01 | 0.004 |
| Colorectal cancer | 329 | 1.03 (1.00, 1.05) | 0.04 | 1.02 (0.99, 1.05) | 0.28 | 1.17 (0.93, 1.47) | 0.18 | 0.23 |
| Pancreatic cancer | 201 | 1.01 (0.97, 1.04) | 0.76 | 1.00 (0.96, 1.04) | 0.98 | 1.04 (0.77, 1.41) | 0.80 | 0.79 |
| Stomach cancer | 105 | 1.07 (1.03, 1.11) | 5.08x10<sup>-04</sup> | 1.06 (1.01, 1.12) | 0.03 | 1.32 (0.84, 2.08) | 0.23 | 0.35 |
| Oesophageal cancer | 226 | 1.06 (1.03, 1.09) | 1.06x10<sup>-04</sup> | 1.07 (1.04, 1.11) | 8.50x10<sup>-05</sup> | 1.20 (0.88, 1.62) | 0.25 | 0.48 |
| Malignant melanoma | 78 | 0.99 (0.94, 1.05) | 0.85 | 0.97 (0.91, 1.04) | 0.40 | 0.91 (0.57, 1.46) | 0.71 | 0.80 |
| Kidney cancer | 137 | 1.09 (1.05, 1.12) | 8.27x10<sup>-07</sup> | 1.08 (1.03, 1.13) | 5.58x10<sup>-04</sup> | 1.08 (0.73, 1.60) | 0.70 | 0.998 |
| Bladder cancer | 78 | 0.98 (0.93, 1.04) | 0.58 | 0.93 (0.87, 1.00) | 0.06 | 0.90 (0.55, 1.45) | 0.66 | 0.88 |
| Brain cancer | 169 | 1.01 (0.97, 1.05) | 0.59 | 0.98 (0.94, 1.03) | 0.44 | 1.29 (0.93, 1.79) | 0.12 | 0.10 |
| Liver cancer | 100 | 1.11 (1.07, 1.15) | 3.18x10<sup>-08</sup> | 1.08 (1.03, 1.13) | 0.003 | 0.81 (0.52, 1.25) | 0.34 | 0.20 |
| Lymphatic cancer | 329 | 1.00 (0.97, 1.03) | 0.91 | 1.01 (0.98, 1.04) | 0.59 | 1.04 (0.82, 1.32) | 0.74 | 0.79 |
| Other cancer | 460 | 0.99 (0.96, 1.01) | 0.22 | 1.00 (0.97, 1.03) | 0.88 | 0.90 (0.74, 1.11) | 0.33 | 0.31 |
| External causes | 206 | 0.97 (0.94, 1.01) | 0.12 | 0.97 (0.93, 1.01) | 0.11 | 1.25 (0.94, 1.67) | 0.13 | 0.08 |
| Other | 696 | 1.00 (0.99, 1.02) | 0.65 | 0.99 (0.97, 1.01) | 0.38 | 1.00 (0.84, 1.18) | 0.97 | 0.94 |

BMI = body mass index; CI = confidence interval; DWH = Durbin-Wu-Hausman; HR = hazard ratio; MR = Mendelian randomization
<sup>1</sup>Number of deaths from all causes or cause-specific mortality
<sup>2</sup>Adjusted for secular trends (date of birth), estimates represent HR with each unit increase in BMI (kg/m<sup>2</sup>)
<sup>3</sup>Additionally adjusted for highest household occupation, education, smoking status, alcohol intake, physical activity and the first ten genetic PCs
<sup>4</sup>P-value for comparing estimates derived from observational and MR analyses using a simplification of the matrix method for DWH test statistic (see Supporting Methods)

**Table 4.**
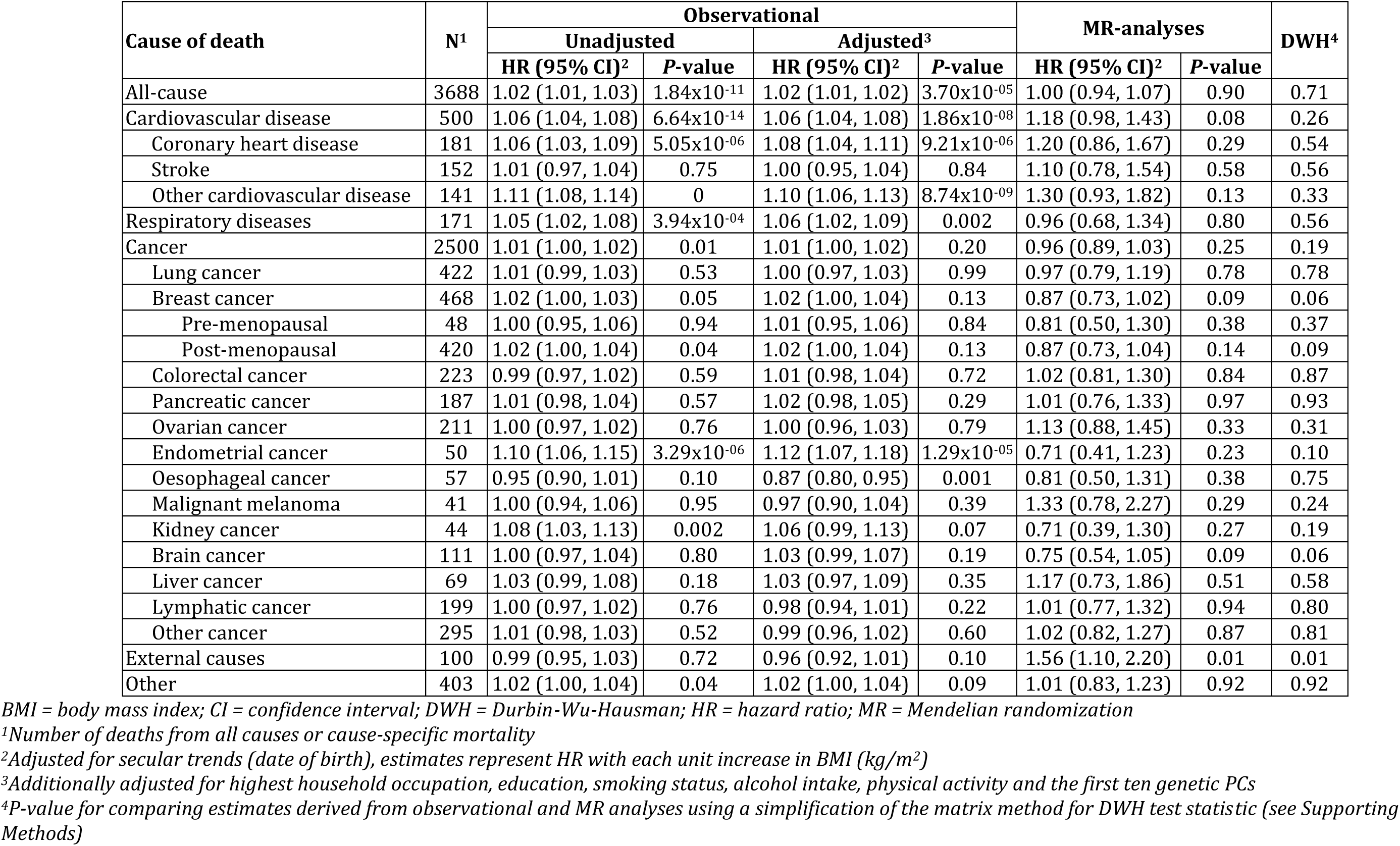
Observational and MR analyses of all-cause and cause-specific mortality by BMI in females of White British ancestry

| Cause of death | N<sup>1</sup> | Observational |  |  |  | MR-analyses |  | DWH<sup>4</sup> |
| --- | --- | --- | --- | --- | --- | --- | --- | --- |
|  |  | Unadjusted |  | Adjusted<sup>3</sup> |  |  |  |  |
|  |  | HR (95% CI)<sup>2</sup> | P-value | HR (95% CI)<sup>2</sup> | P-value | HR (95% CI)<sup>2</sup> | P-value |  |
| All-cause | 3688 | 1.02 (1.01, 1.03) | 1.84x10<sup>-11</sup> | 1.02 (1.01, 1.02) | 3.70x10<sup>-05</sup> | 1.00 (0.94, 1.07) | 0.90 | 0.71 |
| Cardiovascular disease | 500 | 1.06 (1.04, 1.08) | 6.64x10<sup>-14</sup> | 1.06 (1.04, 1.08) | 1.86x10<sup>-08</sup> | 1.18 (0.98, 1.43) | 0.08 | 0.26 |
| Coronary heart disease | 181 | 1.06 (1.03, 1.09) | 5.05x10<sup>-06</sup> | 1.08 (1.04, 1.11) | 9.21x10<sup>-06</sup> | 1.20 (0.86, 1.67) | 0.29 | 0.54 |
| Stroke | 152 | 1.01 (0.97, 1.04) | 0.75 | 1.00 (0.95, 1.04) | 0.84 | 1.10 (0.78, 1.54) | 0.58 | 0.56 |
| Other cardiovascular disease | 141 | 1.11 (1.08, 1.14) | 0 | 1.10 (1.06, 1.13) | 8.74x10<sup>-09</sup> | 1.30 (0.93, 1.82) | 0.13 | 0.33 |
| Respiratory diseases | 171 | 1.05 (1.02, 1.08) | 3.94x10<sup>-04</sup> | 1.06 (1.02, 1.09) | 0.002 | 0.96 (0.68, 1.34) | 0.80 | 0.56 |
| Cancer | 2500 | 1.01 (1.00, 1.02) | 0.01 | 1.01 (1.00, 1.02) | 0.20 | 0.96 (0.89, 1.03) | 0.25 | 0.19 |
| Lung cancer | 422 | 1.01 (0.99, 1.03) | 0.53 | 1.00 (0.97, 1.03) | 0.99 | 0.97 (0.79, 1.19) | 0.78 | 0.78 |
| Breast cancer | 468 | 1.02 (1.00, 1.03) | 0.05 | 1.02 (1.00, 1.04) | 0.13 | 0.87 (0.73, 1.02) | 0.09 | 0.06 |
| Pre-menopausal | 48 | 1.00 (0.95, 1.06) | 0.94 | 1.01 (0.95, 1.06) | 0.84 | 0.81 (0.50, 1.30) | 0.38 | 0.37 |
| Post-menopausal | 420 | 1.02 (1.00, 1.04) | 0.04 | 1.02 (1.00, 1.04) | 0.13 | 0.87 (0.73, 1.04) | 0.14 | 0.09 |
| Colorectal cancer | 223 | 0.99 (0.97, 1.02) | 0.59 | 1.01 (0.98, 1.04) | 0.72 | 1.02 (0.81, 1.30) | 0.84 | 0.87 |
| Pancreatic cancer | 187 | 1.01 (0.98, 1.04) | 0.57 | 1.02 (0.98, 1.05) | 0.29 | 1.01 (0.76, 1.33) | 0.97 | 0.93 |
| Ovarian cancer | 211 | 1.00 (0.97, 1.02) | 0.76 | 1.00 (0.96, 1.03) | 0.79 | 1.13 (0.88, 1.45) | 0.33 | 0.31 |
| Endometrial cancer | 50 | 1.10 (1.06, 1.15) | 3.29x10<sup>-06</sup> | 1.12 (1.07, 1.18) | 1.29x10<sup>-05</sup> | 0.71 (0.41, 1.23) | 0.23 | 0.10 |
| Oesophageal cancer | 57 | 0.95 (0.90, 1.01) | 0.10 | 0.87 (0.80, 0.95) | 0.001 | 0.81 (0.50, 1.31) | 0.38 | 0.75 |
| Malignant melanoma | 41 | 1.00 (0.94, 1.06) | 0.95 | 0.97 (0.90, 1.04) | 0.39 | 1.33 (0.78, 2.27) | 0.29 | 0.24 |
| Kidney cancer | 44 | 1.08 (1.03, 1.13) | 0.002 | 1.06 (0.99, 1.13) | 0.07 | 0.71 (0.39, 1.30) | 0.27 | 0.19 |
| Brain cancer | 111 | 1.00 (0.97, 1.04) | 0.80 | 1.03 (0.99, 1.07) | 0.19 | 0.75 (0.54, 1.05) | 0.09 | 0.06 |
| Liver cancer | 69 | 1.03 (0.99, 1.08) | 0.18 | 1.03 (0.97, 1.09) | 0.35 | 1.17 (0.73, 1.86) | 0.51 | 0.58 |
| Lymphatic cancer | 199 | 1.00 (0.97, 1.02) | 0.76 | 0.98 (0.94, 1.01) | 0.22 | 1.01 (0.77, 1.32) | 0.94 | 0.80 |
| Other cancer | 295 | 1.01 (0.98, 1.03) | 0.52 | 0.99 (0.96, 1.02) | 0.60 | 1.02 (0.82, 1.27) | 0.87 | 0.81 |
| External causes | 100 | 0.99 (0.95, 1.03) | 0.72 | 0.96 (0.92, 1.01) | 0.10 | 1.56 (1.10, 2.20) | 0.01 | 0.01 |
| Other | 403 | 1.02 (1.00, 1.04) | 0.04 | 1.02 (1.00, 1.04) | 0.09 | 1.01 (0.83, 1.23) | 0.92 | 0.92 |

BMI = body mass index; CI = confidence interval; DWH = Durbin-Wu-Hausman; HR = hazard ratio; MR = Mendelian randomization
<sup>1</sup>Number of deaths from all causes or cause-specific mortality
<sup>2</sup>Adjusted for secular trends (date of birth), estimates represent HR with each unit increase in BMI (kg/m<sup>2</sup>)
<sup>3</sup>Additionally adjusted for highest household occupation, education, smoking status, alcohol intake, physical activity and the first ten genetic PCs
<sup>4</sup>P-value for comparing estimates derived from observational and MR analyses using a simplification of the matrix method for DWH test statistic (see Supporting Methods)

In females, adjusted Cox regression analyses provided evidence for an association between higher BMI and increased risk of mortality from endometrial cancer (HR: 1.12; 95% CI: 1.07, 1.18; *P*=1.29×10^−05^), weak evidence for an association of higher BMI with an increased risk of mortality from post-menopausal breast cancer (HR: 1.02; 95% CI: 1.00, 1.04; *P*=0.13) and other causes (HR: 1.02; 95% CI: 1.00, 1.04; *P*=0.09). There was no strong evidence of an association with ovarian cancer (Table 4). There was no strong evidence of an association of BMI with mortality from lung cancer and the estimate of association between higher BMI and mortality from oesophageal cancer was in the opposite direction to that observed in the whole sample (HR: 0.87; 95% CI: 0.80, 0.95; *P*=0.001); however, all CIs overlapped (Table 4).

### Association between the GRS and BMI

Each unit increase in the GRS (comprising 77 SNPs) in the UK Biobank participants of White British ancestry was associated with 0.11kg/m^2^ higher BMI (95% CI: 0.11, 0.11; *P*<1.20×10^−307^), explaining 1.8% of the variance in UK Biobank and was similar between males and females (Table 5).

**Table 5.**
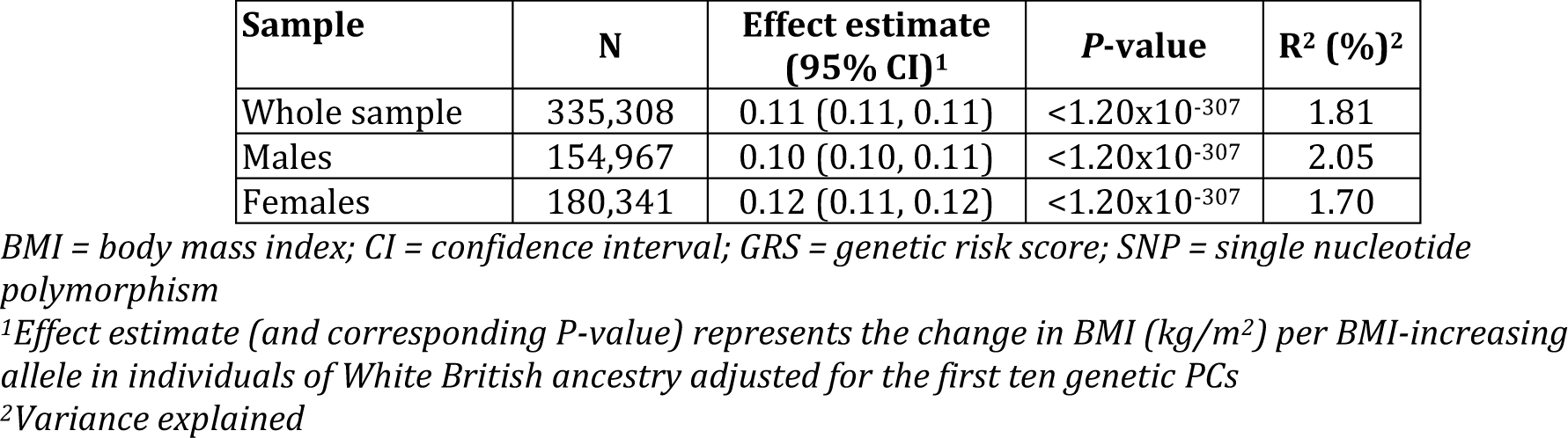
Association between weighted GRS (comprising 77 SNPs) and BMI in UK Biobank participants of White British ancestry

### Confounder analysis

As expected with observational analyses, both BMI and mortality were associated with all covariables including age of initial assessment, sex, smoking status, alcohol consumption, qualifications, employment status and physical activity (Table S4 and 5 for BMI and all-cause mortality, respectively). Unlike the direct measurement of BMI, the GRS was associated with covariables to a much lesser extent, with all estimates near zero (Table S6).

### MR analyses

Within the whole UK Biobank sample, using the GRS as an IV in MR analyses provided estimates of a similar or greater magnitude to observational analyses (albeit, with wider CIs), supporting the causal role of higher BMI in increasing the risk of all-cause mortality (HR: 1.02; 95% CI: 0.97, 1.06; *P*=0.45) and mortality from CVD (HR: 1.12; 95% CI: 1.02, 1.23; *P*=0.02), specifically CHD (HR: 1.19; 95% CI: 1.05, 1.35; *P*=0.01) and those other than stroke and aortic aneurysm (HR: 1.13; 95% CI: 0.93, 1.38; *P*=0.22), as well as mortality from stomach cancer (HR: 1.30; 95% CI: 0.91, 1.86; *P*=0.16) and oesophageal cancer (HR: 1.08; 95% CI: 0.84, 1.38; *P*=0.55) (Table 2). Although CIs were wide, the estimate of the effect of higher BMI on decreasing the risk of mortality from lung cancer was consistent to that obtained in observational analyses (HR: 0.97; 95% CI: 0.84, 1.11; *P*=0.63). There was also evidence supporting the causal role of higher BMI in increasing the risk of mortality from external causes (HR: 1.35; 95% CI: 1.09, 1.67; *P*=0.01), unlike the inverse association obtained in observational analyses (DWH *P*-value for comparison between observational and MR analyses: 0.002). In contrast, the estimates of the effect of higher BMI on mortality from cancer, kidney cancer and liver cancer were attenuated or in the opposite direction, with CIs too wide for comparison to observational analyses or conclusive interpretation (Table 2).

Results for males were similar, in that the estimates of the causal role of higher BMI in increasing the risk of all-cause mortality and mortality from CVDs (including CHD and those other than stroke and aortic aneurysm), stomach cancer, oesophageal cancer and kidney cancer, as well as the decreased risk of mortality from respiratory diseases, lung cancer and bladder cancer, were consistent to or greater than the observational analyses (Table 3). The estimates of the effect of higher BMI on risk of mortality from prostate cancer (HR: 0.73; 95% CI: 0.57, 0.93; *P*=0.01) and external causes (HR: 1.25; 95% CI: 0.94, 1.67; *P*=0.13) were in the opposite direction to those obtained in observational analyses (DWH *P*-value for comparison: 0.004 and 0.08, respectively). The estimates of the effect of higher BMI on mortality from cancer and liver cancer were attenuated or in the opposite direction, with CIs too wide for comparison to observational analyses or conclusive interpretation (Table 3).

In females, the estimates of the effect of higher BMI on increasing the risk of mortality from CVDs (including CHD and those other than stroke) and other causes, as well as the decreased risk of mortality from oesophageal cancer, were consistent to the observational analyses (Table 4). The estimates of the effect of higher BMI on risk of mortality from external causes (HR: 1.56; 95% CI: 1.10, 2.20; *P*=0.01), breast cancer (HR: 0.87; 95% CI: 0.73, 1.02; *P*=0.09), specifically post-menopausal breast cancer (HR: 0.87; 95% CI: 0.73, 1.04; *P*=0.14), and endometrial cancer (HR: 0.71; 95% CI: 0.41, 1.23; *P*=0.23) were in the opposite direction to those obtained in observational analyses (DWH *P*-value for comparison: 0.01, 0.06, 0.09 and 0.10, respectively). Furthermore, the estimates of the effect of higher BMI on mortality from all causes, respiratory diseases, overall cancer and kidney cancer were attenuated or in the opposite direction compared to observational analyses, but with CIs too wide for conclusive interpretation (Table 4).

### Linearity and proportional hazards assumption

The pattern of association between the GRS and all-cause mortality appeared linear (Figure 2); however, the CIs were wide. Assessment of linearity between BMI and most causes of mortality showed a J-shaped association in observational analyses (Figure 3A). When localised average causal effects within strata of instrument-free BMI were used to plot an approximate MR analogue to the plot of mortality against BMI, a J-shape was visible (Figure 3B), but with a smaller value of BMI at which mortality risk was lowest (∼23kg/m^2^ compared to ∼26kg/m^2^ with observational analyses) and apparently flatter over a larger range of BMI, as compared to the observational association. However, CIs were wide with this analysis. Meta-regression testing for a linear trend in the association between the GRS and BMI over quantiles of instrument-free exposure provided some evidence that the GRS-BMI association was non-linear (*P*-value for linear trend = 0.07 and *P*-value for heterogeneity < 0.001). This was primarily driven by the extreme quantiles of BMI, as removal of these quantiles indicated a linear association in the meta-regression (*P*-value for linear trend = 0.99 and *P*-value for heterogeneity < 0.001).

**Figure 2.**
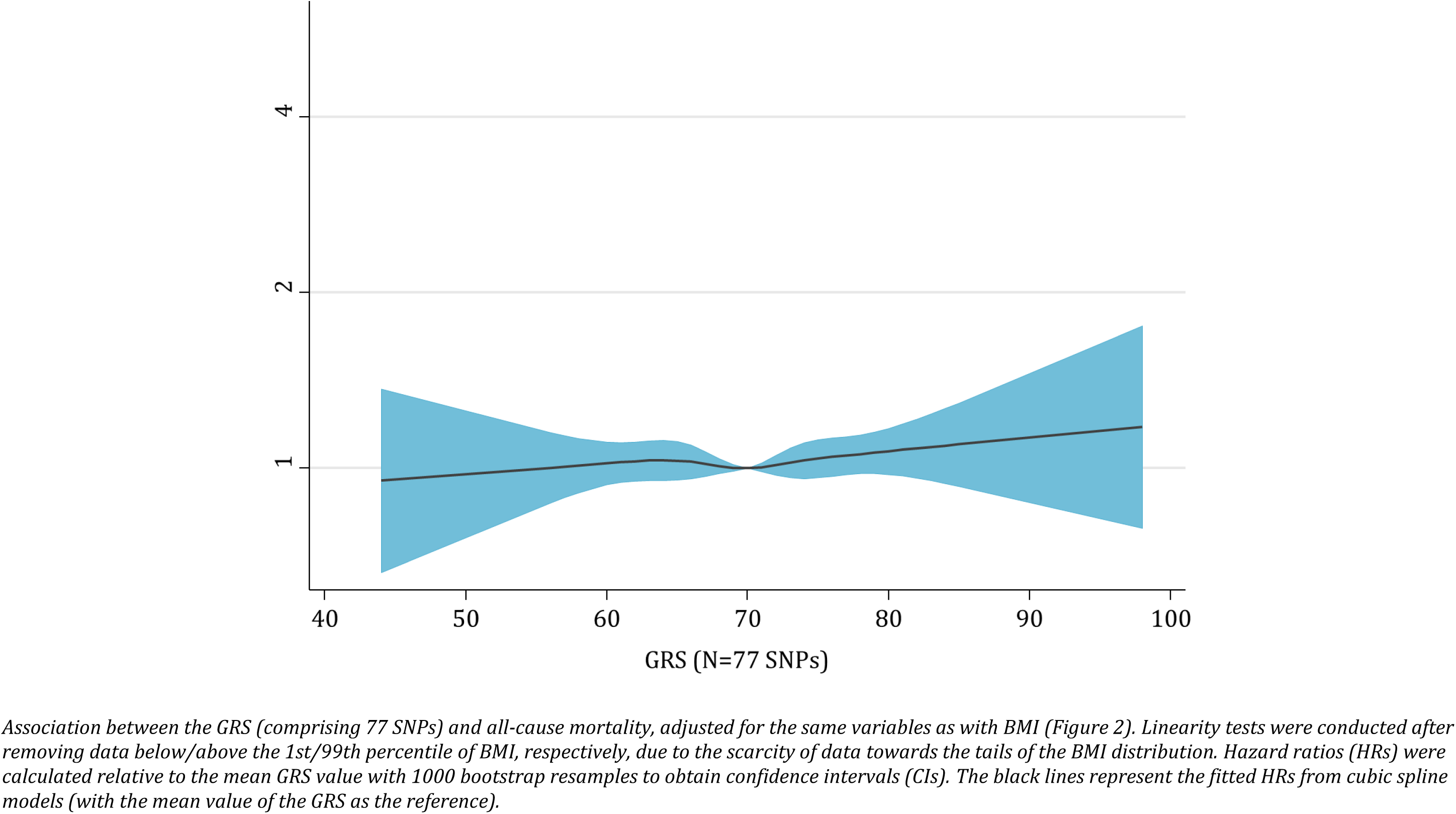
Assessment of linearity in associations of the GRS (comprising 77 SNPs, right) and all-cause mortality in the UK Biobank sample of White British ancestry.

**Figure 3.**
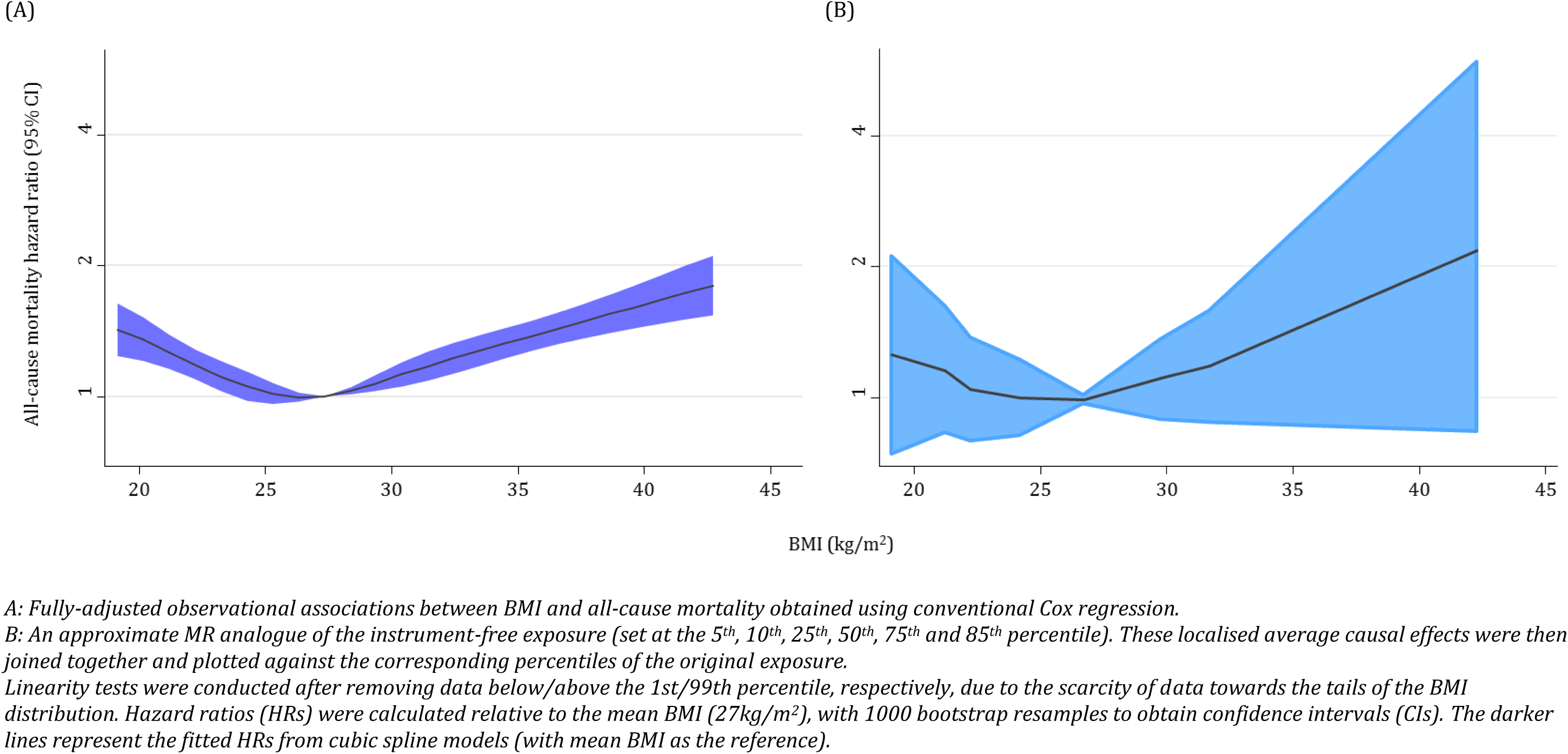
Assessment of linearity in associations of BMI and all-cause mortality in the UK Biobank sample of White British ancestry using BMI (A) and instrument-free BMI (B).

The proportional hazards assumption held for all causes of mortality in both the conventional Cox regression and the MR analyses, especially given the number of tests being performed (Table S7a and S7b for correlation coefficients and corresponding *P*-values for testing the proportional hazards assumption in observational and MR analyses, respectively).

### Sensitivity analyses

Across all methods applied, which assume linearity (including MR-Egger, weighted median- and mode-based estimators as compared to the IVW approach), MR-derived estimates were consistent (Table S8a, S8b and S8c for results in the whole sample, males and females, respectively). Whilst there was no evidence for pleiotropic effects in the estimation of the causal role of BMI in a majority of the mortality causes, the intercept estimate from MR-Egger regression showed marginal evidence for pleiotropy in the association between BMI and mortality from other cancers (intercept: 0.96; 95% CI: 0.93, 1.00; *P*=0.03; HR from MR-Egger regression: 1.25; 95% CI: 0.96, 1.62; *P*=0.09, Table S8a and Figure S1a) and mortality from other causes (intercept: 0.97; 95% CI: 0.94, 0.99; *P*=0.02; HR from MR-Egger regression: 1.27; 95% CI: 1.03, 1.58; *P*=0.03, Figure S1b), suggesting an underestimated MR estimate with negative directional pleiotropy. In males, there was marginal evidence for pleiotropy in the association between BMI and mortality from other cancers (intercept: 0.95; 95% CI: 0.91, 0.99; *P*=0.01; HR from MR-Egger regression: 1.35; 95% CI: 0.97, 1.89; *P*=0.07, Table S8b and Figure S2a) and other causes (intercept: 0.96; 95% CI: 0.93, 1.00; *P*=0.04; HR from MR-Egger regression: 1.27; 95% CI: 0.97, 1.66; *P*=0.08, Figure S2b), similarly suggesting negative directional pleiotropy. There was no strong evidence of directional pleiotropy in female-specific analyses (Table S8c). In both the whole sample and male-specific analyses, the negative directional pleiotropy was likely driven by the rs17024393 SNP (in the association between BMI and mortality from other cancers) and by many SNPs with small positive effects on BMI and small negative effects on mortality (in the association between BMI and mortality from other causes).

Additional adjustment of genotyping chip made no substantive difference to the current MR analyses (Table S9), providing no strong evidence of a differential array effect. When limiting the GRS to exclude the SNPs that may have potentially pleiotropic effects based on previous literature and biological plausibility (N=70), there was no substantive difference in the association between the GRS and BMI (Table S10). The results of the MR analyses were similar to the main analyses (Table S11).

## DISCUSSION

Results supported the causal role of higher BMI in increasing the risk of all-cause mortality and mortality specifically from CVD (including CHD and those other than stroke and aortic aneurysm) plus various cancer sub-types including those of the oesophagus and stomach, as well as decreasing the risk of lung cancer mortality. Sex-stratified analyses were consistent to those in the whole sample and provided additional evidence to support the causal role of higher BMI in increasing the risk of mortality from bladder cancer in males and other causes in females, whilst decreasing the risk of mortality from respiratory diseases in males.

The current results for the most common mortality causes are consistent with previous studies that support the causal role of higher BMI in increasing the risk of all-cause mortality and mortality specifically from vascular diseases and various cancers, whilst decreasing the risk of lung cancer mortality^1, 2, 5, 7, 8, 21^. For example, the largest systematic review and meta-analysis of this relationship (including more than 30 million participants and approximately 3.7 million deaths) showed consistent evidence that each 5kg/m^2^ increment in BMI was associated with a 5% increased risk (95% CI: 4-7%) of all-cause mortality in all participants^21^. Concordant with this, scaling the current results in UK Biobank suggested that each 5kg/m^2^ increase in BMI was associated with a ∼10% increased risk in all-cause mortality. Similarly, and consistent with a collaborative analysis of over 900,000 adults showing an approximate 40% increased risk of vascular mortality with each 5kg/m^2^ higher BMI^1^, scaling the current results to reflect the same increase in BMI implied a ∼75% increased risk of overall CVD and, specifically, more than a 2.5-fold increased risk of CHD.

For cancer, whilst CIs for all MR analyses were wide, the current analyses provided evidence for a causal role of higher BMI increasing the risk of mortality from cancer of the stomach and oesophagus, whilst decreasing the risk of lung cancer mortality in all participants (but CIs were wide). Sex-specific analyses were broadly consistent with an additional inverse association with mortality from respiratory diseases and bladder cancer in males, in whom the effect on kidney cancer mortality was also stronger than in all participants. In females, there was consistent evidence that BMI played a causal role in mortality from other causes. However, these analyses were likely limited by the rarity of mortality from cancer (i.e., many cancers had fewer than 300 cases), accentuated further particularly in analyses within female participants, where many estimates derived from MR analyses were opposite to those from observational analyses or had CIs too wide for interpretation. Despite this, many effect estimates were in the same direction as those derived from previous large-scale meta-analyses and reviews. The association of BMI on incidence of 22 cancer sites in 5.24 million individuals, suggested linear positive relationships with cancers of the kidney, liver, colorectal and ovary and inverse associations with prostate, pre-menopausal breast cancer and lung cancer, the latter being strongly driven by smoking status^25^. Consistent with this, despite estimates from the conventional Cox regression suggesting a positive association between BMI and prostate cancer mortality in UK Biobank, MR analyses provided evidence in the opposite direction (i.e., higher BMI reducing prostate risk). Additionally, in the Million Women Study, incrementally higher BMI was associated with a higher risk of mortality from cancers of the endometrium, oesophagus (specifically, adenocarcinoma subtypes), kidney, pancreas, lymphatic system, ovary, breast cancer (in post-menopausal women) and colorectal cancer (in pre-menopausal women)^5^. However, whilst there was observational evidence for a positive effect on mortality from endometrial cancer and post-menopausal breast cancer, estimates were inverse in MR analyses.

When using instrument-free BMI as the exposure, the association between BMI and all-cause mortality showed a J-shaped pattern but appeared flatter over a larger range of BMI, as compared to the observational association, with a smaller value of BMI at which mortality risk was lowest. Such a difference may be suggestive of confounding in observational associations (i.e., potentially overestimating the harmful effects of underweight whilst underestimating the harmful effects of being overweight and obese); however, we accept wide CIs suggest a need for more power to be conclusive. Likely explanations for a heightened J-shaped curve in observational studies include confounding by lifestyle factors, which may generate a higher risk of mortality in underweight individuals, or reverse causality. Many previous studies used populations comprising older individuals with likely existing illnesses, which may generate a spurious association of lower BMI increasing the risk of mortality (i.e., those who lose weight as a result of disease)^17, 28, 29^. Indeed, in the largest study to date, overestimation of estimates and this characteristic J-shaped association were reported greatest in analyses with the most potential for bias (including all participants, current, former or never smokers and studies with short follow-up durations of <5 years), highlighting the importance for unbiased modes of estimation (such as those used in the current analysis)^21^. Those that attempt to appropriately control for such effects (such as adjusting for baseline traits, restricting analyses to individuals who never smoked or had a longer follow-up), observe an emerging linear association^2, 11, 21, 71, 72^, suggesting an underestimation of the positive link of BMI on mortality but falsely overestimating the effect of low BMI on mortality in a general population. Whilst it is plausible that individuals considered severely and unhealthily underweight have a higher risk of mortality than those within the normal ranges of BMI^73^, the current findings in this large population of healthy individuals support a more linear association, with lower BMI being protective. Furthermore, the lowest risk of mortality occurred at approximately 23kg/m^2^ when using MR methodology, as opposed to being overweight (i.e., a BMI of 25-30kg/m^2^), which was observed in the current observational analyses and has been implied previously by some of the existing observational literature^13^. Therefore, a stable BMI within the ‘normal’ range (i.e., 18.5-25kg/m^2^) may be the most beneficially healthy with regards to reducing mortality risk, with any reduction within that range likely to be beneficial^8, 21^.

Use of the UK Biobank study has enabled an MR investigation of the causal role of BMI in all-cause and cause-specific mortality. This has been afforded given the large sample size, comprehensive genetic and phenotypic measurements and detailed death registry information. Whilst the power to detect associations with MR analyses was low for many mortality causes in the current study, given the incidence of the outcomes tested, this will increase in the coming years where incidence and mortality from many causes will approximately double by 2022^57^.

The MR concept rests on several key assumptions^31–34^: (i) the IV (here, the GRS) must be associated with the exposure (here, BMI); (ii) the IV must be independent of the factors that act as confounders of the association between the exposure and the outcome (here, all-cause and cause-specific mortality); and (iii) there must be no independent pathway between the IV and outcome other than through the exposure – horizontal pleiotropy^34^. These assumptions were tested where possible, but it can be difficult to directly assess whether possible sources of pleiotropy are horizontal (independent of the exposure) or vertical (involved in the causal pathway between the exposure and outcome)^66^. Within the context of the current study, sensitivity analyses conducted provided little evidence of pleiotropy in MR estimates and awarded greater confidence in the validity of the instrument used and, thus, MR-derived estimates. Reverse causality is an important source of bias in observational estimates of the association between BMI and mortality and may be the driver of the characteristic J-shaped association. However, whilst it is possible that mortality from specific causes can influence the relative distribution of genetic variants within a selected sample^74^, it is likely that this potential bias is less marked than that seen in observational studies (especially in light of the use of common genetic variants to predict a complex risk factor like BMI).

The UK Biobank study is comprised of relatively healthy volunteer participants and is a unique opportunity to undertake these analyses. Even in this study, however, the sample sizes available were limited. With this, it is likely that participants with higher education and socioeconomic position and different causes of death and estimates of association may not be representative of and generalisable to the wider national or international populations. Furthermore, to remove any possible effect of genetic confounding, the current analyses were restricted to those only of White British ancestry, limiting the generalisability of results to other ancestral groups. Generally, the current analyses were likely limited by the rarity of many mortality causes (e.g., the number of individuals who had died from diabetes and other cardiovascular-related traits were insufficient to include), accentuated further in sex-stratified analyses. Therefore, the potential role of chance and low statistical power in many relatively weak associations such as those observed between BMI and mortality from the various rare mortality causes cannot be ruled out.

### Conclusions

This study represents the application of MR to assess the causal effect of higher BMI on the risk and cause of mortality. Results supported the causal role of higher BMI in increasing the risk of all-cause mortality and mortality specifically from CVD, various cancers and other causes. Along with the comprehensive studies with greater numbers of deaths in combination with the application of robust causal inference methods such as those employed here that appropriately account for the heavy burden of confounding, reverse causation and bias within observational epidemiological designs, our results further highlight the need for a global effort to reduce the rising population trends for excess weight from an early age.

## Acknowledgements

The authors are grateful to the UK Biobank participants. This research has been conducted using the UK Biobank Resource under the Application Number 16391.

