## Supplementary Materials for "Body mass index and mortality in UK Biobank: revised estimates using Mendelian randomization"

**Corresponding author:**

Dr Kaitlin H Wade

Oakfield House

Oakfield Road

Clifton

Bristol

BS8 2BN

### Supporting Methods

*Standard exclusions*

After preliminary exclusions, 484,514 individuals had a valid measure of body mass index (BMI), plausible age and death data, along with available genetic data. The following exclusions were made to the dataset required for survival analyses prior to all analyses based on in-house quality control (QC) parameters (total excluded = 149,206; see figure below). Individuals who were outliers in heterozygosity and missing rates (f.22027) were already excluded from imputed data.

- Sex mismatch (N=367) – derived by comparing the genetic sex variable (f.22001), as determined by Affymetrix, with reported sex (f.31) of the participant.

- Sex-chromosome aneuploidy (N=643) – individuals with sex chromosome karyotypes putatively different from XX or XY (f.22019). Of these, 177 individuals overlapped with the sex-mismatch list (above).
- Relatedness – minimally related individuals were removed (N=79,034), which were defined as the first individual in a related pair (3rd degree or closer) based on an algorithm applied to the list of all the related pairs provided by UK Biobank. This number also included any individuals who appeared to be highly related to a very large number (>200) of individuals, derived using the list of individuals excluded from the kinship inference (N=9). Of these, 106 individuals overlapped with those excluded based on sex mismatch and sex-chromosome aneuploidy (above). Once removed, the remaining subset was the maximal set of unrelated individuals in UK Biobank.
- Ancestry – we used stringent criteria for excluding those not of White British ancestry, defined as those who self-reported as "White" and "British" and had very similar genetic ancestry based on a principal components analysis of the genotypes (N=77,722). Of these, 8,277 individuals overlapped with those excluded based on sex mismatch, sex-chromosome aneuploidy and relatedness (above).

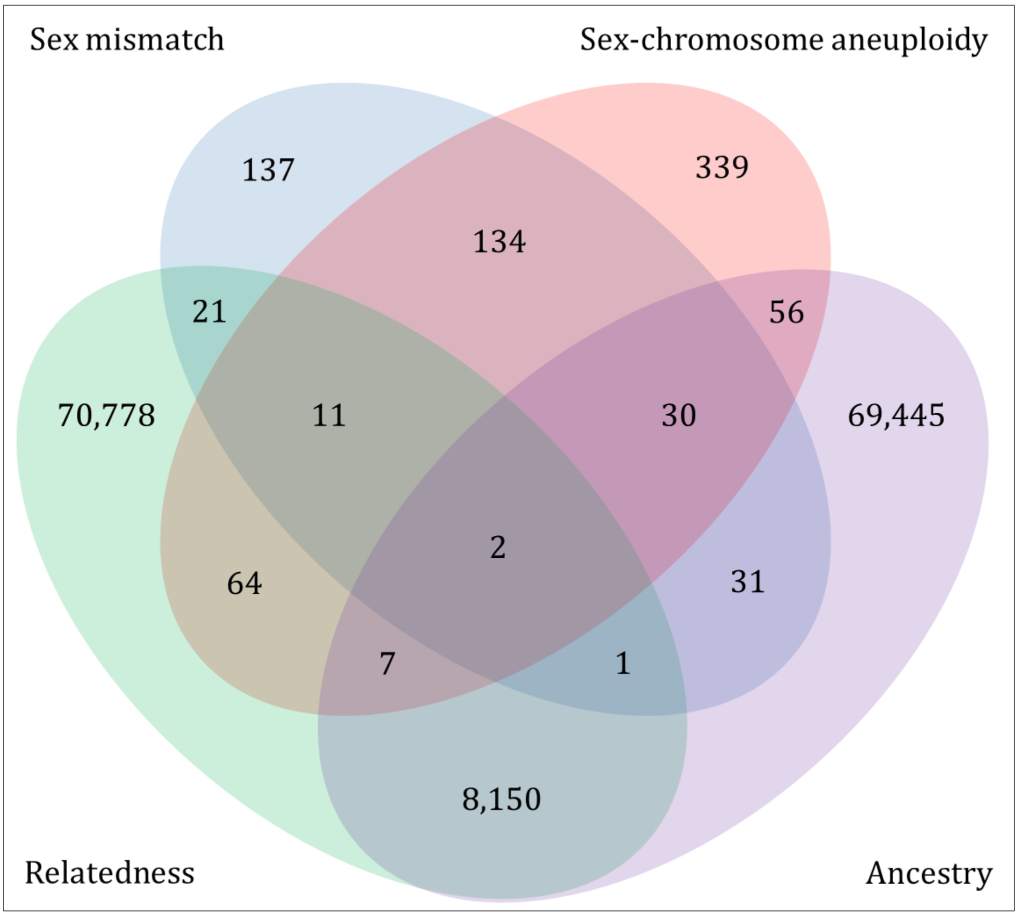

*Test for endogeneity between conventional Cox regression and MR analyses*

A simplification of the matrix method for the Durbin-Wu-Hausman (DWH) test for endogeneity was used to compare effect estimates derived from conventional Cox regression and Mendelian randomization (MR) analyses. For one instrumental variable (here, a weighted genetic risk score [GRS]), the test statistic can be simplified to the following formula:

$$Z = \frac{(\beta_{CCR} - \beta_{MR})^2}{({SE_{CCR}}^2 - {SE_{MR}}^2)}$$

where $\beta_{CCR}$ and $\beta_{MR}$ are the effect estimates obtained from the conventional Cox regression and MR analyses, respectively, and $SE_{CCR}$ and $SE_{MR}$ are the corresponding standard errors. The test statistics, $Z$, has a chi-squared distribution with one degree of freedom.

Table S1a. Descriptive statistics for UK Biobank mortality data

| Cause of death | ICD10 codes | Total number of deaths in UK Biobank | | |
|---|---|---|---|---|
| | | Whole sample | With valid data | White British |
| All-cause | All | 14,417 | 11,161 | 9.570 |
| Cardiovascular disease[1] | I*; G459 | 2,999 | 2,332 | 1.967 |
| Coronary heart disease[2] | I209-I259; I516 | 1,661 | 1,291 | 1.087 |
| Stroke[3] | I600-I698; G459 | 550 | 428 | 346 |
| Aortic aneurysm | I710-I719 | 152 | 120 | 109 |
| Other cardiovascular diseases | All other I | 636 | 493 | 425 |
| **Diabetes[4]** | **E10-E149** | **70** | **50** | **37** |
| Respiratory diseases[5] | J* | 834 | 631 | 532 |
| Cancer[6] | C* | 8,286 | 6,484 | 5.613 |
| Lung cancer | C33-C349 | 1,507 | 1,160 | 993 |
| Breast cancer | C50-C509 | 739 | 560 | 472 |
| Prostate cancer | C61-C619 | 440 | 348 | 308 |
| Colorectal cancer | C180-C219 | 822 | 646 | 552 |
| Pancreatic cancer | C250-C259 | 556 | 446 | 388 |
| Stomach cancer | C160-C169 | 201 | 162 | 144 |
| Ovarian cancer | C56-C570 | 306 | 247 | 211 |
| Endometrial cancer | C54-C549 | 76 | 56 | 50 |
| **Gallbladder cancer** | **C23-C249** | **55** | **41** | **37** |
| Oesophageal cancer | C15-C159 | 389 | 310 | 283 |
| Malignant melanoma | C43-C449 | 173 | 131 | 119 |
| **Thyroid cancer** | **C73-C739** | **16** | **11** | **9** |
| Kidney cancer | C64-C669 | 245 | 196 | 181 |
| Bladder cancer | C67-C679 | 161 | 122 | 101 |
| Brain cancer | C71-C729 | 408 | 322 | 280 |
| Liver cancer | C220-C229 | 266 | 208 | 169 |
| **Cervical cancer** | **C53-C539** | **16** | **13** | **12** |
| **Uterine cancer** | **C55-C559** | **40** | **29** | **21** |
| Lymphatic cancer | C810-C964 | 730 | 592 | 528 |
| Other cancers | All other C | 1140 | 884 | 755 |
| **Kidney disease** | **N00-N299** | **34** | **23** | **16** |
| External causes[7] | V*; W*; X*; Y* | 496 | 359 | 306 |
| Other[8] | - | 1,698 | 1,282 | 1.099 |

*ICD = International Classification of Diseases*
*Mortality causes in bold were not included in analyses due to small number of deaths (<40): cancer of the gallbladder (N=37), thyroid (N=9), cervix (N=12) and uterus (N=21), and kidney disease (N=16).*
**Any code beginning with the indicated letter*
[1]*Cardiovascular disease consisted of all disease of the circulatory system listed in ICD 10, including coronary heart disease and stroke.*
[2]*Coronary heart disease is a narrowing of the arteries supplying the heart muscle and may be considered synonymous with ischemic heart disease or coronary artery disease.*
[3]*Stroke included bleeding from (haemorrhagic stroke) or blockage of (ischemic stroke) the arteries supplying the brain, as well as transient ischemic attacks ("mini-strokes").*
[4]*Diabetes included insulin-dependent and non-insulin-dependent diabetes mellitus.*
[5]*Respiratory diseases included all non-neoplasmic diseases of the lungs, pleura and respiratory tract.*
[6]*Cancers excluded benign or in-situ neoplasms.*
[7]*External causes consisted of accidents and violence, including suicide and conditions consequent to accidents and violence*
[8]*Other includes all other causes of mortality that aren't listed.*

Table S1b. Descriptive statistics for UK Biobank mortality data stratified by sex

| **Cause of death** | **ICD10 codes** | **Total number of deaths in UK Biobank (with valid data for main analyses)** | |
|---|---|---|---|
| | | **Males** | **Females** |
| All-cause | All | 5,882 | 3,688 |
| Cardiovascular disease[1] | I*; G459 | 1,467 | 500 |
| Coronary heart disease[2] | I209-I259; I516 | 906 | 181 |
| Stroke[3] | I600-I698; G459 | 194 | 152 |
| **Aortic aneurysm** | **I710-I719** | 83 | **26** |
| Other cardiovascular diseases | All other I | 284 | 141 |
| **Diabetes[4]** | **E10-E149** | **29** | **8** |
| Respiratory diseases[5] | J* | 361 | 171 |
| Cancer[6] | C* | 3,113 | 2,500 |
| Lung cancer | C33-C349 | 571 | 422 |
| **Breast cancer** | **C50-C509** | **4** | 468 |
| Prostate cancer | C61-C619 | 308 | - |
| Colorectal cancer | C180-C219 | 329 | 223 |
| Pancreatic cancer | C250-C259 | 201 | 187 |
| **Stomach cancer** | **C160-C169** | 105 | **39** |
| Ovarian cancer | C56-C570 | - | 211 |
| Endometrial cancer | C54-C549 | - | 50 |
| **Gallbladder cancer** | **C23-C249** | **16** | **21** |
| Oesophageal cancer | C15-C159 | 226 | 57 |
| Malignant melanoma | C43-C449 | 78 | 41 |
| **Thyroid cancer** | **C73-C739** | **2** | **7** |
| Kidney cancer | C64-C669 | 137 | 44 |
| **Bladder cancer** | **C67-C679** | 78 | **23** |
| Brain cancer | C71-C729 | 169 | 111 |
| Liver cancer | C220-C229 | 100 | 69 |
| **Cervical cancer** | **C53-C539** | **-** | **12** |
| **Uterine cancer** | **C55-C559** | **-** | **21** |
| Lymphatic cancer | C810-C964 | 329 | 199 |
| Other cancers | All other C | 460 | 295 |
| **Kidney disease** | **N00-N299** | **10** | **6** |
| External causes[7] | V*; W*; X*; Y* | 206 | 100 |
| Other[8] | - | 696 | 403 |

*ICD = International Classification of Diseases*

*In addition to the mortality causes with a collectively small number of deaths in the whole sample (Table S1a), the sex-specific mortality causes in bold were also not included in analyses due to small numbers of deaths (<40) when stratified by sex: aortic aneurysm in females (N=26), diabetes (N=29/8 in males/females, respectively), breast cancer in males (N=4) and cancer of the stomach in females (N=39), gallbladder (N=16/21 in males/females, respectively) and bladder in females (N=23).*

**Any code beginning with the indicated letter*

[1]*Cardiovascular disease consisted of all disease of the circulatory system listed in ICD 10, including coronary heart disease and stroke.*

[2]*Coronary heart disease is a narrowing of the arteries supplying the heart muscle and may be considered synonymous with ischemic heart disease or coronary artery disease.*

[3]*Stroke included bleeding from (haemorrhagic stroke) or blockage of (ischemic stroke) the arteries supplying the brain, as well as transient ischemic attacks ("mini-strokes").*

[4]*Diabetes included insulin-dependent and non-insulin-dependent diabetes mellitus.*

[5]*Respiratory diseases included all non-neoplasmic diseases of the lungs, pleura and respiratory tract.*

[6]*Cancers excluded benign or in-situ neoplasms.*

[7]*External causes consisted of accidents and violence, including suicide and conditions consequent to accidents and violence*

[8]*Other includes all other causes of mortality that aren't listed.*

Table S2. Genetic variants associated with BMI available in UK Biobank individuals

| Genetic variant | Gene | Chr | bp | Effect allele[1] | Other allele[1] | EAF[2] | Beta (SE)[3] | P-value[3] | Imputation quality[2] |
|---|---|---|---|---|---|---|---|---|---|
| rs1000940 | *RABEP1* | 17 | 5283252 | G | A | 0.30 | 0.07 (0.01) | $1.45x10^{-07}$ | 0.998 |
| rs10132280 | *STXBP6* | 14 | 25928179 | C | A | 0.70 | 0.11 (0.01) | $2.53x10^{-17}$ | 0.988 |
| rs1016287 | *LINC01122* | 2 | 59305625 | T | C | 0.30 | 0.10 (0.01) | $1.03x10^{-14}$ | 0.997 |
| rs10182181 | *ADCY3* | 2 | 25150296 | G | A | 0.49 | 0.17 (0.01) | $1.42x10^{-48}$ | 0.996 |
| rs10733682 | *LMX1B* | 9 | 129460914 | A | G | 0.47 | 0.06 (0.01) | $1.63x10^{-07}$ | 0.962 |
| rs10938397 | *GNPDA2* | 4 | 45182527 | G | A | 0.43 | 0.14 (0.01) | $8.89x10^{-34}$ | 1.000 |
| rs10968576 | *LINGO2* | 9 | 28414339 | G | A | 0.31 | 0.12 (0.01) | $7.87x10^{-23}$ | 1.000 |
| rs11030104 | *BDNF* | 11 | 27684517 | A | G | 0.80 | 0.18 (0.01) | $1.74x10^{-35}$ | 0.999 |
| rs11057405 | *CLIP1* | 12 | 122781897 | G | A | 0.90 | 0.14 (0.02) | $3.09x10^{-14}$ | 1.000 |
| rs11126666 | *KCNK3* | 2 | 26928811 | A | G | 0.26 | 0.02 (0.01) | 0.16 | 0.995 |
| rs11165643 | *PTBP2* | 1 | 96924097 | T | C | 0.58 | 0.08 (0.01) | $9.45x10^{-13}$ | 0.998 |
| rs11191560 | *NT5C2* | 10 | 104869038 | C | T | 0.08 | 0.12 (0.02) | $1.42x10^{-08}$ | 0.999 |
| rs11583200 | *ELAVL4* | 1 | 50559820 | C | T | 0.40 | 0.07 (0.01) | $1.78x10^{-09}$ | 0.993 |
| rs1167827 | *HIP1* | 7 | 75163169 | G | A | 0.57 | 0.11 (0.01) | $6.77x10^{-20}$ | 1.000 |
| rs11688816 | *EHBP1* | 2 | 63053048 | G | A | 0.54 | 0.06 (0.01) | $1.76x10^{-07}$ | 0.993 |
| rs11727676 | *HHIP* | 4 | 145659064 | T | C | 0.91 | 0.04 (0.02) | 0.03 | 1.000 |
| rs11847697 | *PRKD1* | 14 | 30515112 | T | C | 0.05 | 0.12 (0.03) | $3.99x10^{-05}$ | 1.000 |
| rs12286929 | *CADM1* | 11 | 115022404 | G | A | 0.52 | 0.08 (0.01) | $9.77x10^{-11}$ | 0.996 |
| rs12401738 | *FUBP1* | 1 | 78446761 | A | G | 0.33 | 0.07 (0.01) | $4.54x10^{-10}$ | 0.995 |
| rs12429545 | *OLFM4* | 13 | 54102206 | A | G | 0.13 | 0.13 (0.02) | $3.37x10^{-13}$ | 0.979 |
| rs12446632 | *GPRC5B* | 16 | 19935389 | G | A | 0.86 | 0.13 (0.02) | $1.77x10^{-15}$ | 1.000 |
| rs12566985 | *FPGT-TNNI3K* | 1 | 75002193 | G | A | 0.45 | 0.07 (0.01) | $1.14x10^{-09}$ | 0.998 |
| rs12885454 | *PRKD1* | 14 | 29736838 | C | A | 0.65 | 0.07 (0.01) | $1.82x10^{-09}$ | 0.998 |
| rs12940622 | *RPTOR* | 17 | 78615571 | G | A | 0.56 | 0.08 (0.01) | $2.82x10^{-12}$ | 0.999 |
| rs13021737 | *TMEM18* | 2 | 632348 | G | A | 0.83 | 0.25 (0.02) | $6.91x10^{-59}$ | 1.000 |
| rs13078960 | *CADM2* | 3 | 85807590 | G | T | 0.20 | 0.10 (0.01) | $4.95x10^{-11}$ | 0.994 |
| rs13107325 | *SLC39A8* | 4 | 103188709 | T | C | 0.07 | 0.25 (0.02) | $2.79x10^{-29}$ | 1.000 |
| rs13191362 | *PARK2* | 6 | 163033350 | A | G | 0.88 | 0.10 (0.02) | $7.78x10^{-09}$ | 0.995 |
| rs13201877 | *IFNGR1* | 6 | 137675541 | G | A | 0.13 | 0.04 (0.02) | 0.01 | 0.978 |
| rs1441264 | *MIR548A2* | 13 | 79580919 | A | G | 0.60 | 0.10 (0.01) | $7.82x10^{-17}$ | 0.953 |
| rs1460676 | *FIGN* | 2 | 164567689 | C | T | 0.16 | 0.07 (0.02) | $3.94x10^{-05}$ | 0.996 |
| rs1516725 | *ETV5* | 3 | 185824004 | C | T | 0.86 | 0.16 (0.02) | $6.60x10^{-22}$ | 0.992 |
| rs1528435 | *UBE2E3* | 2 | 181550962 | T | C | 0.62 | 0.08 (0.01) | $4.57x10^{-11}$ | 0.997 |

| | | | | | | | | | |
|---|---|---|---|---|---|---|---|---|---|
| rs1558902 | *FTO* | 16 | 53803574 | A | T | 0.39 | 0.36 (0.01) | $5.55x10^{-201}$ | 1.000 |
| rs16851483 | *RASA2* | 3 | 141275436 | T | G | 0.07 | 0.17 (0.02) | $3.35x10^{-13}$ | 0.998 |
| rs16907751 | *ZBTB10* | 8 | 81375457 | C | T | 0.90 | 0.10 (0.02) | $1.15x10^{-06}$ | 0.991 |
| rs16951275 | *MAP2K5* | 15 | 68077168 | T | C | 0.77 | 0.14 (0.01) | $9.56x10^{-23}$ | 0.999 |
| rs17001654 | *SCARB2* | 4 | 77129568 | G | C | 0.15 | 0.09 (0.02) | $2.65x10^{-07}$ | 0.973 |
| rs17024393 | *GNAT2* | 1 | 110154688 | C | T | 0.03 | 0.31 (0.04) | $1.15x10^{-17}$ | 0.998 |
| rs17094222 | *HIF1AN* | 10 | 102395440 | C | T | 0.21 | 0.07 (0.01) | $4.00x10^{-07}$ | 0.991 |
| rs17203016 | *CREB1* | 2 | 208255518 | G | A | 0.20 | 0.07 (0.01) | $4.19x10^{-07}$ | 0.992 |
| rs17405819 | *HNF4G* | 8 | 76806584 | T | C | 0.71 | 0.09 (0.01) | $9.06x10^{-14}$ | 1.000 |
| rs17724992 | *PGPEP1* | 19 | 18454825 | A | G | 0.73 | 0.08 (0.01) | $2.93x10^{-09}$ | 0.991 |
| rs1808579 | *C18orf8* | 18 | 21104888 | C | T | 0.52 | 0.10 (0.01) | $3.89x10^{-18}$ | 0.998 |
| rs1928295 | *TLR4* | 9 | 120378483 | T | C | 0.57 | 0.06 (0.01) | $1.49x10^{-06}$ | 1.000 |
| rs2033529 | *TDRG1* | 6 | 40348653 | G | A | 0.28 | 0.11 (0.01) | $3.37x10^{-17}$ | 0.992 |
| rs2033732 | *RALYL* | 8 | 85079709 | C | T | 0.75 | 0.05 (0.01) | $9.75x10^{-05}$ | 1.000 |
| rs205262 | *C6orf106* | 6 | 34563164 | G | A | 0.27 | 0.14 (0.01) | $1.35x10^{-27}$ | 0.998 |
| rs2075650 | *TOMM40* | 19 | 45395619 | A | G | 0.85 | 0.09 (0.02) | $1.39x10^{-07}$ | 1.000 |
| rs2080454 | *CBLN1* | 16 | 49062590 | C | A | 0.39 | 0.05 (0.01) | $1.88x10^{-05}$ | 0.989 |
| rs2112347 | *POC5* | 5 | 75015242 | T | G | 0.63 | 0.14 (0.01) | $2.19x10^{-29}$ | 1.000 |
| rs2121279 | *LRP1B* | 2 | 143043285 | T | C | 0.12 | 0.06 (0.02) | 0.002 | 0.991 |
| rs2176040 | *LOC646736* | 2 | 227092802 | A | G | 0.35 | 0.02 (0.01) | 0.13 | 1.000 |
| rs2176598 | *HSD17B12* | 11 | 43864278 | T | C | 0.25 | 0.09 (0.01) | $4.91x10^{-12}$ | 1.000 |
| rs2207139 | *TFAP2B* | 6 | 50845490 | G | A | 0.17 | 0.20 (0.02) | $9.26x10^{-37}$ | 1.000 |
| rs2245368 | *PMS2L11* | 7 | 76608143 | C | T | 0.18 | 0.11 (0.02) | $1.30x10^{-13}$ | 1.000 |
| rs2287019 | *QPCTL* | 19 | 46202172 | C | T | 0.82 | 0.15 (0.02) | $1.35x10^{-24}$ | 0.984 |
| rs2365389 | *FHIT* | 3 | 61236462 | C | T | 0.58 | 0.08 (0.01) | $2.93x10^{-11}$ | 0.994 |
| rs2650492 | *SBK1* | 16 | 28333411 | A | G | 0.29 | 0.09 (0.01) | $8.09x10^{-12}$ | 0.988 |
| rs2820292 | *NAV1* | 1 | 201784287 | C | A | 0.56 | 0.10 (0.01) | $9.47x10^{-17}$ | 1.000 |
| rs2836754 | *ETS2* | 21 | 40291740 | C | T | 0.62 | 0.06 (0.01) | $3.73x10^{-07}$ | 1.000 |
| rs29941 | *KCTD15* | 19 | 34309532 | G | A | 0.67 | 0.08 (0.01) | $9.00x10^{-10}$ | 1.000 |
| rs3101336 | *NEGR1* | 1 | 72751185 | C | T | 0.61 | 0.10 (0.01) | $1.09x10^{-18}$ | 1.000 |
| rs3736485 | *DMXL2* | 15 | 51748610 | A | G | 0.47 | 0.06 (0.01) | $6.18x10^{-07}$ | 0.995 |
| rs3810291 | *ZC3H4* | 19 | 47569003 | A | G | 0.66 | 0.13 (0.01) | $2.84x10^{-26}$ | 1.000 |
| rs3817334 | *MTCH2* | 11 | 47650993 | T | C | 0.40 | 0.12 (0.01) | $8.05x10^{-23}$ | 1.000 |
| rs3849570 | *GBE1* | 3 | 81792112 | A | C | 0.35 | 0.06 (0.01) | $2.36x10^{-06}$ | 1.000 |
| rs3888190 | *ATP2A1* | 16 | 28889486 | A | C | 0.39 | 0.13 (0.01) | $7.32x10^{-28}$ | 1.000 |
| rs4256980 | *TRIM66* | 11 | 8673939 | G | C | 0.65 | 0.08 (0.01) | $8.94x10^{-12}$ | 0.996 |

| | | | | | | | | | |
|---|---|---|---|---|---|---|---|---|---|
| rs4740619 | *C9orf93* | 9 | 15634326 | T | C | 0.55 | 0.09 (0.01) | $3.61x10^{-14}$ | 0.999 |
| rs4771122* | *MTIF3* | 13 | 28020180 | G | A | 0.22 | 0.04 (0.01) | 0.01 | 1.000 |
| rs4787491 | *INO80E* | 16 | 30015337 | G | A | 0.53 | 0.10 (0.01) | $2.61x10^{-16}$ | 0.998 |
| rs492400 | *USP37* | 2 | 219349752 | C | T | 0.43 | 0.06 (0.01) | $2.10x10^{-06}$ | 0.996 |
| rs543874 | *SEC16B* | 1 | 177889480 | G | A | 0.20 | 0.23 (0.01) | $2.11x10^{-58}$ | 1.000 |
| rs6091540 | *ZFP64* | 20 | 51087862 | C | T | 0.70 | 0.09 (0.01) | $3.81x10^{-12}$ | 0.995 |
| rs6465468 | *ASB4* | 7 | 95169514 | T | G | 0.30 | 0.03 (0.01) | 0.02 | 0.990 |
| rs6477694 | *EPB41L4B* | 9 | 111932342 | C | T | 0.36 | 0.06 (0.01) | $1.80x10^{-06}$ | 0.990 |
| rs6567160 | *MC4R* | 18 | 57829135 | C | T | 0.23 | 0.25 (0.01) | $6.70x10^{-75}$ | 0.998 |
| rs657452 | *AGBL4* | 1 | 49589847 | A | G | 0.40 | 0.08 (0.01) | $1.88x10^{-11}$ | 0.987 |
| rs6804842 | *RARB* | 3 | 25106437 | G | A | 0.57 | 0.05 (0.01) | $4.39x10^{-06}$ | 0.991 |
| rs7138803 | *BCDIN3D* | 12 | 50247468 | A | G | 0.36 | 0.13 (0.01) | $1.93x10^{-28}$ | 1.000 |
| rs7141420 | *NRXN3* | 14 | 79899454 | T | C | 0.52 | 0.10 (0.01) | $1.05x10^{-17}$ | 0.985 |
| rs7164727 | *LOC100287559* | 15 | 73093991 | T | C | 0.66 | 0.09 (0.01) | $9.25x10^{-14}$ | 0.995 |
| rs7239883 | *LOC284260* | 18 | 40147671 | G | A | 0.38 | 0.05 (0.01) | $7.87x10^{-06}$ | 0.994 |
| rs7243357 | *GRP* | 18 | 56883319 | T | G | 0.82 | 0.09 (0.02) | $9.41x10^{-09}$ | 0.990 |
| rs758747 | *NLRC3* | 16 | 3627358 | T | C | 0.29 | 0.06 (0.01) | $9.10x10^{-06}$ | 0.978 |
| rs7599312 | *ERBB4* | 2 | 213413231 | G | A | 0.73 | 0.08 (0.01) | $7.12x10^{-09}$ | 0.979 |
| rs7715256 | *GALNT10* | 5 | 153537893 | G | T | 0.43 | 0.08 (0.01) | $4.76x10^{-11}$ | 0.999 |
| rs7899106 | *GRID1* | 10 | 87410904 | G | A | 0.05 | 0.13 (0.03) | $1.68x10^{-06}$ | 0.987 |
| rs7903146 | *TCF7L2* | 10 | 114758349 | C | T | 0.71 | 0.08 (0.01) | $1.04x10^{-10}$ | 1.000 |
| rs9374842 | *LOC285762* | 6 | 120185665 | T | C | 0.77 | 0.05 (0.01) | $8.20x10^{-04}$ | 0.997 |
| rs9400239 | *FOXO3* | 6 | 108977663 | C | T | 0.69 | 0.08 (0.01) | $3.48x10^{-10}$ | 0.994 |
| rs9540493 | *MIR548X2* | 13 | 66205704 | A | G | 0.46 | 0.07 (0.01) | $9.03x10^{-09}$ | 0.989 |
| rs9641123 | *CALCR* | 7 | 93197732 | C | G | 0.40 | 0.04 (0.01) | $2.65x10^{-04}$ | 0.996 |
| rs977747 | *TAL1* | 1 | 47684677 | T | G | 0.43 | 0.08 (0.01) | $2.29x10^{-11}$ | 0.997 |
| rs9914578 | *SMG6* | 17 | 2005136 | G | C | 0.21 | 0.05 (0.01) | $8.34x10^{-04}$ | 0.995 |
| rs9925964 | *KAT8* | 16 | 31129895 | A | G | 0.65 | 0.12 (0.01) | $1.95x10^{-24}$ | 0.998 |

*BMI = body mass index; bp = base pair; Chr = chromosome; EAF = effect allele frequency; SE = standard error*

*[1]Effect and other alleles associated with an increasing BMI, according to the most recent genome-wide association study of BMI[2]*

*[2]Minor allele frequency (MAF) of each genetic variant in UK Biobank and corresponding imputation quality (the latter of which was based on the whole of UK Biobank)*

*[3]Beta and corresponding standard error (SE) and P-value represent the change in BMI ($kg/m^2$) per BMI-increasing allele of each genetic variant in individuals of White British ancestry adjusted for the first ten genetic principal components.*

**The rs4771122 SNP served as the closest proxy (with an $r^2$=0.876, distance=2398bp), according to SNAP (http://archive.broadinstitute.org/mpg/snap/ldsearch.php) for the rs12016871 on chromosome 13, which was not available in the UK Biobank genetic data.*

Table S3. Genetic variants excluded in from sensitivity analyses

| SNP | Gene | Chromosome | Reason for exclusion |
|---|---|---|---|
| rs977747 | *TAL1* | 1 | All ancestries |
| rs1460676 | *FIGN* | 2 | All ancestries |
| rs17203016 | *CREB1* | 2 | All ancestries |
| rs2176040 | *LOC646736* | 2 | European men |
| rs492400 | *USP37* | 2 | European men |
| rs13107325 | *SLC39A8* | 4 | Pleiotropic effects |
| rs17001654 | *SCARB2* | 4 | Pleiotropic effects |
| rs7715256 | *GALNT10* | 5 | All ancestries |
| rs13201877 | *IFNGR1* | 6 | All ancestries |
| rs9374842 | *LOC285762* | 6 | European population based |
| rs1167827 | *HIP1* | 7 | Pleiotropic effects |
| rs6465468 | *ASB4* | 7 | European women |
| rs9641123 | *CALCR* | 7 | European population based |
| rs16907751 | *ZBTB10* | 8 | European men |
| rs7903146 | *TCF7L2* | 10 | Identified in Corbin *et al.*[3] |
| rs11030104 | *BDNF* | 11 | Pleiotropic effects |
| rs1441264 | *MIR548A2* | 13 | All ancestries |
| rs9540493 | *MIR548X2* | 13 | European population based |
| rs7164727 | *LOC100287559* | 15 | All ancestries |
| rs2080454 | *CBLN1* | 16 | All ancestries |
| rs3888190 | *ATP2A1* | 16 | Pleiotropic effects |
| rs4787491 | *INO80E* | 16 | European population based |
| rs9914578 | *SMG6* | 17 | All ancestries |
| rs7239883 | *LOC284260* | 18 | European women |
| rs2075650 | *TOMM40* | 19 | Pleiotropic effects |
| rs6091540 | *ZFP64* | 20 | European women |
| rs2836754 | *ETS2* | 21 | All ancestries |

*Consistent with previous studies[3-5], these genetic variants were excluded in sensitivity analyses to test the robustness of the GRS used in MR analyses*

Table S4. Association of BMI with participants' characteristics

| **Variable** | **N** | **Estimate (95% CI)[1]** | ***P*-value** |
|---|---|---|---|
| Age (years) | 335,308 | 0.03 (0.03, 0.03) | $3.82x10^{-182}$ |
| Sex (% of males) | 335,308 | 0.81 (0.78, 0.84) | $<1.20x10^{-307}$ |
| *Smoking status* | 334,142 | | |
| Never | | reference | |
| Former | | 0.81 (0.78, 0.84) | $<1.20x10^{-307}$ |
| Current | | -0.07 (-0.12, -0.01) | 0.02 |
| *Alcohol drinker status* | 335,074 | | |
| Never | | reference | |
| Former | | 0.27 (0.14, 0.40) | $2.74x10^{-05}$ |
| Current | | -0.69 (-0.78, -0.59) | $1.21x10^{-47}$ |
| *Highest qualifications* | 275,544 | | |
| College or University degree | | reference | |
| A-levels | | 0.59 (0.54, 0.65) | $2.97x10^{-102}$ |
| O-levels | | 1.01 (0.96, 1.05) | $<1.20x10^{-307}$ |
| CSEs | | 1.49 (1.42, 1.57) | $<1.20x10^{-307}$ |
| NVQ/HND/HNC | | 1.70 (1.64, 1.77) | $<1.20x10^{-307}$ |
| Other professional qualifications | | 1.16 (1.09, 1.24) | $1.20x10^{-203}$ |
| *Current employment status* | 332,835 | | |
| In paid employment or self-employed | | reference | |
| Retired | | 0.28 (0.25, 0.31) | $2.07x10^{-57}$ |
| Looking after home/family | | -0.70 (-0.80, -0.60) | $2.35x10^{-41}$ |
| Unable to work due to sickness/disability | | 2.46 (2.37, 2.56) | $<1.20x10^{-307}$ |
| Unemployed | | 0.83 (0.69, 0.97) | $5.64x10^{-31}$ |
| Doing unpaid or voluntary work | | -0.73 (-0.98, -0.48) | $8.62x10^{-09}$ |
| Full or part-time student | | -0.77 (-1.14, -0.40) | $5.02x10^{-05}$ |
| Days/week spent doing vigorous physical activity | 319,813 | -0.24 (-0.24, -0.23) | $<1.20x10^{-307}$ |
| Genotyping chip[2] | 335,308 | 0.60 (0.55, 0.66) | $3.30x10^{-100}$ |

*BMI = body mass index; CI = confidence interval; CSE = certificate of secondary education; HNC = higher national certificate; HND = higher national diploma; NVQ = national vocational qualification*

[1]*Estimates represent the difference in BMI (kg/m$^2$) per unit increase in each continuous, categorical or binary variable in individuals of White British ancestry adjusted for the first ten genetic principal components.*

[2]*There was evidence of differential array effect on markers scattered across the genome; therefore, the UK BiLEVE study genotyped on the Affymetrix Axiom Array was considered as a confounder*

Table S5. Association of participants' characteristics and all-cause mortality

| Variable | HR (95% CI)[1] | *P*-value |
|---|---|---|
| Age (years) | 0.95 (0.94, 0.96) | $5.00x10^{-19}$ |
| Sex (% of males) | 1.80 (1.72, 1.87) | $1.08x10^{-170}$ |
| *Smoking status* | | |
| Never | reference | |
| Former | 1.51 (1.45, 1.58) | $1.18x10^{-71}$ |
| Current | 3.22 (3.05, 3.41) | $<1.20x10^{-307}$ |
| *Alcohol drinker status* | | |
| Never | reference | |
| Former | 1.87 (1.65, 2.13) | $1.07x10^{-21}$ |
| Current | 0.93 (0.84, 1.03) | 0.15 |
| *Highest qualifications* | | |
| College or University degree | reference | |
| A-levels | 1.10 (1.02, 1.19) | 0.02 |
| O-levels | 1.10 (1.04, 1.17) | 0.002 |
| CSEs | 1.35 (1.20, 1.51) | $3.01x10^{-07}$ |
| NVQ/HND/HNC | 1.34 (1.23, 1.46) | $6.67x10^{-12}$ |
| Other professional qualifications | 1.12 (1.02, 1.23) | 0.02 |
| *Current employment status* | | |
| In paid employment or self-employed | reference | |
| Retired | 1.12 (1.06, 1.18) | $9.41x10^{-05}$ |
| Looking after home/family | 1.04 (0.87, 1.24) | 0.67 |
| Unable to work due to sickness/disability | 5.11 (4.73, 5.51) | $<1.20x10^{-307}$ |
| Unemployed | 2.29 (1.93, 2.70) | $4.04x10^{-22}$ |
| Doing unpaid or voluntary work | 0.85 (0.58, 1.24) | 0.40 |
| Full or part-time student | 1.92 (1.11, 3.31) | 0.02 |
| Days/week spent doing vigorous physical activity | 0.94 (0.93, 0.95) | $3.28x10^{-29}$ |
| Genotyping chip[2] | 1.33 (1.25, 1.42) | $4.85x10^{-20}$ |

*BMI = body mass index; CI = confidence interval; CSE = certificate of secondary education; HNC = higher national certificate; HND = higher national diploma; NVQ = national vocational qualification*

[1]*Estimates represent the difference in hazards for all-cause mortality per unit increase in each continuous, categorical or binary variable in individuals of White British ancestry adjusted for the first ten genetic principal components.*

[2]*There was evidence of differential array effect on markers scattered across the genome; therefore, the UK BiLEVE study genotyped on the Affymetrix Axiom Array was considered as a confounder*

Table S6. Characteristics of participants according to the BMI GRS (comprising 77 SNPs)

| Variable | N | Estimate (95% CI)[1] | *P*-value |
|---|---|---|---|
| Age (years) | 335,308 | -0.005 (-0.01, 0.0004) | 0.07 |
| Sex (% of males) | 335,308 | 0.0003 (-0.00003, 0.001) | 0.08 |
| Smoking status | 334,142 | 0.002 (0.001, 0.002) | $5.53x10^{-19}$ |
| Alcohol drinker status | 335,074 | -0.0004 (-0.001, -0.0002) | $2.33x10^{-04}$ |
| Highest qualifications | 332,835 | 0.001 (0.0002, 0.001) | 0.01 |
| Current employment status | 275,544 | 0.003 (0.002, 0.004) | $1.30x10^{-08}$ |
| Days/week spent doing vigorous physical activity | 319,813 | 0.001 (0.0003, 0.003) | 0.02 |
| Genotyping chip[2] | 335,308 | 0.0004 (0.0002, 0.001) | $3.01x10^{-06}$ |

*BMI = body mass index; CI = confidence interval; CSE = certificate of secondary education; GRS = genetic risk score; HNC = higher national certificate; HND = higher national diploma; NVQ = national vocational qualification; SNP = single nucleotide polymorphism*

[1]*Estimates represent the difference in each variable with each unit increase in the GRS (comprising 77 SNPs) in individuals of White British ancestry adjusted for the first ten genetic principal components.*

[2]*There was evidence of differential array effect on markers scattered across the genome; therefore, the UK BiLEVE study genotyped on the Affymetrix Axiom Array was considered as a confounder*

Table S7a. Statistical test of the proportional hazards assumption in conventional Cox regression

| **Cause of death** | **Correlation coefficient (and corresponding *P*-values) for the test of proportional hazards[1]** | | |
|---|---|---|---|
| | **Whole sample** | **Males** | **Females** |
| All-cause | -0.01 (0.64) | 0.001 (0.94) | -0.02 (0.34) |
| Cardiovascular disease | -0.08 (0.004) | -0.06 (0.06) | -0.14 (0.03) |
| Coronary heart disease | -0.01 (0.76) | 0.01 (0.82) | -0.09 (0.42) |
| Stroke | 0.01 (0.85) | 0.06 (0.56) | -0.04 (0.70) |
| Aortic aneurysm | -0.28 (0.02) | -0.31 (0.02) | - |
| Other cardiovascular diseases | -0.18 (0.003) | -0.17 (0.01) | -0.19 (0.09) |
| Respiratory diseases | -0.05 (0.41) | 0.02 (0.83) | -0.05 (0.66) |
| Cancer | 0.01 (0.42) | 0.02 (0.33) | 0.001 (0.98) |
| Lung cancer | 0.06 (0.19) | 0.17 (0.002) | -0.07 (0.29) |
| Breast cancer | - | - | 0.10 (0.07) |
| Pre-menopausal | - | - | 0.24 (0.12) |
| Post-menopausal | - | - | 0.11 (0.05) |
| Prostate cancer | - | -0.03 (0.69) | - |
| Colorectal cancer | -0.03 (0.49) | -0.04 (0.58) | -0.04 (0.62) |
| Pancreatic cancer | -0.11 (0.08) | -0.09 (0.30) | -0.13 (0.16) |
| Stomach cancer | 0.01 (0.91) | 0.08 (0.51) | - |
| Ovarian cancer | - | - | -0.11 (0.16) |
| Endometrial cancer | - | - | 0.22 (0.21) |
| Oesophageal cancer | -0.08 (0.26) | -0.10 (0.24) | 0.04 (0.78) |
| Malignant melanoma | -0.11 (0.29) | -0.08 (0.56) | -0.15 (0.40) |
| Kidney cancer | 0.08 (0.38) | 0.08 (0.49) | 0.12 (0.55) |
| Bladder cancer | -0.10 (0.44) | -0.07 (0.60) | - |
| Brain cancer | 0.05 (0.50) | 0.004 (0.97) | 0.11 (0.30) |
| Liver cancer | -0.05 (0.63) | -0.05 (0.67) | -0.11 (0.48) |
| Lymphatic cancer | -0.03 (0.62) | -0.02 (0.79) | -0.06 (0.50) |
| Other cancer | 0.09 (0.04) | 0.08 (0.17) | 0.11 (0.13) |
| External causes | 0.09 (0.18) | 0.08 (0.30) | 0.09 (0.41) |
| Other | -0.04 (0.29) | -0.02 (0.71) | -0.07 (0.23) |

[1]*Values represent the correlation coefficient (and corresponding P-values) between the first scaled Schoenfeld residual (obtained from the fully adjusted survival analysis of BMI on each mortality outcome) in conventional Cox regression and the natural logarithm of follow-up time (age). Positive/negative values suggest that the relationship between BMI and mortality increases/decreases over the follow-up period, respectively.*

Table S7b. Statistical test of the proportional hazards assumption using Mendelian randomization

| **Cause of death** | **Correlation coefficient (and corresponding *P*-values) for the test of proportional hazards[1]** | | |
|---|---|---|---|
| | **Whole sample** | **Males** | **Females** |
| All-cause | 0.01 (0.26) | 0.01 (0.44) | 0.01 (0.47) |
| Cardiovascular disease | -0.02 (0.39) | 0.01 (0.80) | -0.11 (0.06) |
| Coronary heart disease | -0.04 (0.31) | -0.01 (0.82) | -0.19 (0.07) |
| Stroke | 0.04 (0.57) | 0.17 (0.07) | -0.07 (0.50) |
| Aortic aneurysm | 0.11 (0.35) | 0.14 (0.29) | - |
| Other cardiovascular diseases | -0.05 (0.44) | -0.03 (0.71) | -0.07 (0.52) |
| Respiratory diseases | -0.02 (0.79) | 0.01 (0.87) | -0.05 (0.68) |
| Cancer | 0.03 (0.04) | 0.02 (0.28) | 0.04 (0.08) |
| Lung cancer | 0.004 (0.92) | 0.03 (0.63) | -0.03 (0.61) |
| Breast cancer | - | - | 0.04 (0.42) |
| Pre-menopausal | - | - | 0.02 (0.88) |
| Post-menopausal | - | - | 0.05 (0.40) |
| Prostate cancer | - | 0.12 (0.08) | - |
| Colorectal cancer | 0.10 (0.05) | 0.03 (0.59) | 0.17 (0.02) |
| Pancreatic cancer | 0.08 (0.17) | 0.07 (0.42) | 0.10 (0.25) |
| Stomach cancer | 0.05 (0.67) | 0.09 (0.49) | - |
| Ovarian cancer | - | - | -0.12 (0.14) |
| Endometrial cancer | - | - | 0.20 (0.26) |
| Oesophageal cancer | -0.03 (0.64) | 0.01 (0.92) | -0.14 (0.37) |
| Malignant melanoma | 0.18 (0.09) | 0.23 (0.08) | 0.08 (0.66) |
| Kidney cancer | -0.01 (0.88) | -0.04 (0.74) | 0.06 (0.78) |
| Bladder cancer | 0.19 (0.12) | 0.20 (0.14) | - |
| Brain cancer | -0.01 (0.93) | -0.09 (0.31) | 0.15 (0.17) |
| Liver cancer | -0.14 (0.15) | -0.20 (0.11) | 0.01 (0.93) |
| Lymphatic cancer | -0.03 (0.60) | 0.004 (0.96) | -0.07 (0.39) |
| Other cancer | 0.09 (0.05) | 0.07 (0.22) | 0.12 (0.11) |
| External causes | 0.09 (0.19) | 0.05 (0.50) | 0.17 (0.13) |
| Other | 0.01 (0.84) | 0.01 (0.85) | 0.003 (0.96) |

[1]*Values represent the correlation coefficient (and corresponding P-values) between the first scaled Schoenfeld residual (obtained from the survival analysis of BMI on each mortality outcome) in Mendelian randomization analyses and the natural logarithm of follow-up time (age). Positive/negative values suggest that the relationship between BMI and mortality increases/decreases over the follow-up period, respectively.*

Table S8a. MR-Egger analysis of BMI on all-cause and cause-specific mortality in UK Biobank participants of White British ancestry

| **Cause of death** | **IVW** | | **Intercept of MR-Egger** | | **MR-Egger** | | **Weighted median** | | **Weighted mode** | |
|---|---|---|---|---|---|---|---|---|---|---|
| | **HR (95% CI)[1]** | ***P*** | **HR (95% CI)[1]** | ***P*** | **HR (95% CI)[1]** | ***P*** | **HR (95% CI)[1]** | ***P*** | **HR (95% CI)[1]** | ***P*** |
| All-cause | 1.02 (0.99, 1.06) | 0.25 | 1.00 (0.99, 1.01) | 0.76 | 1.03 (0.95, 1.13) | 0.46 | 1.04 (0.98, 1.08) | 0.30 | 1.05 (0.99, 1.11) | 0.14 |
| Cardiovascular disease | 1.07 (1.00, 1.15) | 0.05 | 1.00 (0.98, 1.02) | 0.87 | 1.06 (0.89, 1.25) | 0.52 | 1.08 (0.96, 1.17) | 0.25 | 1.21 (1.04, 1.41) | 0.02 |
| Coronary heart disease | 1.09 (0.99, 1.19) | 0.08 | 1.00 (0.97, 1.03) | 0.81 | 1.12 (0.88, 1.41) | 0.35 | 1.03 (0.94, 1.24) | 0.31 | 1.06 (0.89, 1.28) | 0.50 |
| Stroke | 0.98 (0.84, 1.15) | 0.83 | 1.02 (0.97, 1.08) | 0.41 | 0.84 (0.57, 1.26) | 0.40 | 0.92 (0.72, 1.14) | 0.40 | 0.82 (0.59, 1.14) | 0.24 |
| Aortic aneurysm | 0.86 (0.64, 1.16) | 0.32 | 1.05 (0.95, 1.15) | 0.33 | 0.61 (0.29, 1.30) | 0.20 | 0.80 (0.50, 1.23) | 0.29 | 0.82 (0.46, 1.44) | 0.48 |
| Other cardiovascular diseases | 1.18 (1.02, 1.35) | 0.02 | 0.99 (0.94, 1.03) | 0.53 | 1.30 (0.92, 1.83) | 0.13 | 1.18 (0.96, 1.51) | 0.14 | 1.21 (0.89, 1.65) | 0.22 |
| Respiratory diseases | 1.03 (0.90, 1.18) | 0.69 | 1.02 (0.98, 1.07) | 0.33 | 0.88 (0.63, 1.24) | 0.46 | 0.95 (0.79, 1.18) | 0.70 | 0.89 (0.67, 1.19) | 0.44 |
| Cancer | 0.99 (0.95, 1.03) | 0.69 | 1.00 (0.99, 1.01) | 0.99 | 0.99 (0.90, 1.10) | 0.88 | 1.00 (0.94, 1.07) | 0.90 | 1.01 (0.93, 1.09) | 0.80 |
| Lung cancer | 0.97 (0.89, 1.06) | 0.51 | 1.02 (0.99, 1.05) | 0.24 | 0.86 (0.68, 1.07) | 0.18 | 0.95 (0.82, 1.11) | 0.51 | 0.94 (0.77, 1.14) | 0.52 |
| Colorectal cancer | 1.05 (0.91, 1.20) | 0.49 | 1.02 (0.97, 1.06) | 0.47 | 0.94 (0.67, 1.32) | 0.70 | 1.07 (0.87, 1.29) | 0.57 | 1.09 (0.86, 1.39) | 0.47 |
| Pancreatic cancer | 1.07 (0.92, 1.25) | 0.35 | 1.03 (0.98, 1.08) | 0.25 | 0.88 (0.60, 1.28) | 0.50 | 1.00 (0.82, 1.31) | 0.82 | 1.02 (0.77, 1.34) | 0.91 |
| Stomach cancer | 1.15 (0.91, 1.46) | 0.25 | 0.94 (0.87, 1.02) | 0.13 | 1.73 (0.96, 3.10) | 0.07 | 1.57 (0.93, 1.96) | 0.11 | 1.71 (1.00, 2.90) | 0.05 |
| Oesophageal cancer | 1.17 (0.98, 1.39) | 0.08 | 0.99 (0.93, 1.05) | 0.67 | 1.27 (0.83, 1.97) | 0.27 | 1.01 (0.84, 1.46) | 0.50 | 1.00 (0.69, 1.46) | 0.996 |
| Malignant melanoma | 1.15 (0.88, 1.51) | 0.31 | 1.04 (0.95, 1.13) | 0.37 | 0.87 (0.44, 1.71) | 0.68 | 0.89 (0.68, 1.57) | 0.90 | 1.02 (0.62, 1.69) | 0.93 |
| Kidney cancer | 0.95 (0.77, 1.18) | 0.65 | 1.03 (0.96, 1.10) | 0.45 | 0.79 (0.46, 1.35) | 0.38 | 1.05 (0.67, 1.37) | 0.83 | 0.92 (0.56, 1.50) | 0.73 |
| Bladder cancer | 0.84 (0.63, 1.12) | 0.23 | 0.95 (0.87, 1.05) | 0.31 | 1.17 (0.58, 2.37) | 0.66 | 0.90 (0.57, 1.44) | 0.72 | 1.09 (0.61, 1.95) | 0.78 |
| Brain cancer | 1.02 (0.85, 1.23) | 0.80 | 1.01 (0.95, 1.07) | 0.76 | 0.96 (0.61, 1.52) | 0.85 | 1.11 (0.80, 1.47) | 0.58 | 1.16 (0.77, 1.77) | 0.48 |
| Liver cancer | 1.00 (0.80, 1.25) | 0.98 | 1.03 (0.96, 1.10) | 0.46 | 0.82 (0.47, 1.44) | 0.49 | 0.95 (0.66, 1.36) | 0.72 | 1.08 (0.69, 1.70) | 0.74 |
| Lymphatic cancer | 1.03 (0.91, 1.17) | 0.62 | 1.00 (0.96, 1.04) | 0.98 | 1.03 (0.75, 1.40) | 0.86 | 1.04 (0.85, 1.27) | 0.66 | 0.99 (0.76, 1.29) | 0.95 |
| Other cancers | 0.96 (0.87, 1.07) | 0.47 | 0.96 (0.93, 1.00) | 0.03 | 1.25 (0.96, 1.62) | 0.09 | 1.00 (0.89, 1.20) | 0.69 | 1.04 (0.85, 1.27) | 0.70 |
| External causes | 1.22 (1.02, 1.46) | 0.03 | 1.04 (0.98, 1.10) | 0.21 | 0.95 (0.61, 1.47) | 0.80 | 1.07 (0.83, 1.44) | 0.51 | 1.06 (0.76, 1.48) | 0.74 |
| Other | 1.01 (0.92, 1.10) | 0.88 | 0.97 (0.94, 0.99) | 0.02 | 1.27 (1.03, 1.58) | 0.03 | 1.07 (0.97, 1.26) | 0.13 | 1.21 (1.02, 1.43) | 0.03 |

*BMI = body mass index; CI = confidence interval; HR = hazard ratio; IVW = inverse variance weighted; MR = Mendelian randomization*

[1]*Adjusted for secular trends (date of birth), sex and the first ten genetic principal components, estimates represent HR with each unit increase in BMI ($kg/m^2$)*

Table S8b. MR-Egger analysis of BMI on all-cause and cause-specific mortality in UK Biobank males of White British ancestry

| **Cause of death** | **IVW** | | **Intercept of MR-Egger** | | **MR-Egger** | | **Weighted median** | | **Weighted mode** | |
|---|---|---|---|---|---|---|---|---|---|---|
| | **HR (95% CI)[1]** | ***P*** | **HR (95% CI)[1]** | ***P*** | **HR (95% CI)[1]** | ***P*** | **HR (95% CI)[1]** | ***P*** | **HR (95% CI)[1]** | ***P*** |
| All-cause | 1.02 (0.98, 1.06) | 0.28 | 0.99 (0.98, 1.00) | 0.13 | 1.09 (0.99, 1.20) | 0.07 | 1.02 (0.97, 1.11) | 0.29 | 1.03 (0.95, 1.12) | 0.42 |
| Cardiovascular disease | 1.07 (0.99, 1.15) | 0.11 | 0.99 (0.97, 1.02) | 0.54 | 1.12 (0.93, 1.36) | 0.23 | 1.04 (0.93, 1.19) | 0.43 | 1.04 (0.88, 1.23) | 0.65 |
| Coronary heart disease | 1.09 (0.98, 1.21) | 0.11 | 0.99 (0.96, 1.02) | 0.51 | 1.18 (0.91, 1.53) | 0.21 | 1.07 (0.94, 1.28) | 0.32 | 1.10 (0.90, 1.35) | 0.35 |
| Stroke | 1.00 (0.81, 1.23) | 0.98 | 1.00 (0.94, 1.07) | 0.92 | 0.98 (0.58, 1.65) | 0.93 | 0.98 (0.70, 1.26) | 0.71 | 0.91 (0.59, 1.40) | 0.66 |
| Aortic aneurysm | 0.86 (0.62, 1.21) | 0.39 | 1.06 (0.95, 1.18) | 0.29 | 0.57 (0.25, 1.33) | 0.19 | 0.66 (0.48, 1.33) | 0.37 | 0.71 (0.38, 1.32) | 0.28 |
| Other cardiovascular diseases | 1.11 (0.93, 1.33) | 0.24 | 0.97 (0.92, 1.03) | 0.35 | 1.35 (0.86, 2.11) | 0.19 | 1.14 (0.86, 1.52) | 0.29 | 1.14 (0.81, 1.62) | 0.45 |
| Respiratory diseases | 1.03 (0.88, 1.20) | 0.69 | 1.02 (0.97, 1.07) | 0.52 | 0.92 (0.63, 1.35) | 0.67 | 1.07 (0.77, 1.23) | 0.83 | 0.87 (0.62, 1.22) | 0.42 |
| Cancer | 1.00 (0.95, 1.06) | 0.98 | 0.99 (0.97, 1.01) | 0.30 | 1.07 (0.93, 1.23) | 0.33 | 1.03 (0.94, 1.12) | 0.58 | 1.02 (0.91, 1.15) | 0.69 |
| Lung cancer | 0.94 (0.83, 1.06) | 0.32 | 1.01 (0.97, 1.05) | 0.52 | 0.86 (0.63, 1.17) | 0.32 | 0.89 (0.77, 1.13) | 0.47 | 0.85 (0.65, 1.11) | 0.24 |
| Prostate cancer | 0.91 (0.76, 1.09) | 0.32 | 0.96 (0.91, 1.02) | 0.20 | 1.19 (0.76, 1.86) | 0.43 | 1.01 (0.76, 1.25) | 0.81 | 1.05 (0.76, 1.45) | 0.77 |
| Colorectal cancer | 1.07 (0.91, 1.26) | 0.40 | 1.01 (0.96, 1.06) | 0.69 | 0.99 (0.66, 1.49) | 0.98 | 1.00 (0.81, 1.34) | 0.82 | 1.04 (0.73, 1.49) | 0.82 |
| Pancreatic cancer | 1.14 (0.93, 1.39) | 0.21 | 1.02 (0.96, 1.09) | 0.45 | 0.96 (0.58, 1.58) | 0.86 | 0.96 (0.76, 1.44) | 0.80 | 0.98 (0.66, 1.45) | 0.93 |
| Stomach cancer | 1.12 (0.84, 1.50) | 0.42 | 0.98 (0.89, 1.07) | 0.63 | 1.32 (0.64, 2.70) | 0.45 | 1.45 (0.77, 2.04) | 0.33 | 1.62 (0.86, 3.06) | 0.14 |
| Oesophageal cancer | 1.21 (0.99, 1.48) | 0.06 | 0.97 (0.91, 1.04) | 0.43 | 1.45 (0.89, 2.38) | 0.14 | 1.17 (0.86, 1.62) | 0.31 | 1.33 (0.85, 2.09) | 0.22 |
| Malignant melanoma[2] | 1.03 (0.74, 1.43) | 0.85 | 1.02 (0.92, 1.13) | 0.73 | 0.91 (0.40, 2.05) | 0.81 | 1.14 (0.64, 1.76) | 0.76 | 1.14 (0.61, 2.11) | 0.68 |
| Kidney cancer | 1.04 (0.79, 1.36) | 0.77 | 1.02 (0.94, 1.11) | 0.66 | 0.91 (0.46, 1.78) | 0.77 | 1.06 (0.65, 1.47) | 0.98 | 0.92 (0.55, 1.57) | 0.77 |
| Bladder cancer | 0.80 (0.58, 1.10) | 0.17 | 0.93 (0.84, 1.03) | 0.15 | 1.36 (0.61, 3.06) | 0.45 | 0.90 (0.53, 1.53) | 0.68 | 1.04 (0.54, 1.99) | 0.91 |
| Brain cancer | 1.12 (0.90, 1.40) | 0.29 | 1.01 (0.95, 1.09) | 0.69 | 1.02 (0.59, 1.75) | 0.96 | 1.16 (0.74, 1.66) | 0.58 | 1.33 (0.79, 2.21) | 0.28 |
| Liver cancer | 1.04 (0.78, 1.39) | 0.78 | 0.98 (0.89, 1.07) | 0.65 | 1.21 (0.59, 2.47) | 0.60 | 1.29 (0.76, 1.86) | 0.46 | 1.35 (0.75, 2.44) | 0.32 |
| Lymphatic cancer | 1.02 (0.87, 1.19) | 0.80 | 0.99 (0.94, 1.04) | 0.61 | 1.12 (0.76, 1.65) | 0.57 | 1.02 (0.86, 1.39) | 0.45 | 1.18 (0.85, 1.63) | 0.32 |
| Other cancers | 0.92 (0.80, 1.05) | 0.22 | 0.95 (0.91, 0.99) | 0.01 | 1.35 (0.97, 1.89) | 0.07 | 1.06 (0.83, 1.28) | 0.81 | 1.08 (0.84, 1.38) | 0.57 |
| External causes | 1.09 (0.88, 1.34) | 0.42 | 1.05 (0.98, 1.12) | 0.16 | 0.78 (0.47, 1.30) | 0.34 | 0.90 (0.71, 1.32) | 0.84 | 0.92 (0.62, 1.36) | 0.67 |
| Other | 0.98 (0.88, 1.09) | 0.70 | 0.96 (0.93, 1.00) | 0.04 | 1.27 (0.97, 1.66) | 0.08 | 1.09 (0.93, 1.29) | 0.31 | 1.16 (0.93, 1.44) | 0.19 |

*BMI = body mass index; CI = confidence interval; HR = hazard ratio; IVW = inverse variance weighted; MR = Mendelian randomization*

*[1]Adjusted for secular trends (date of birth) and the first ten genetic principal components, estimates represent HR with each unit increase in BMI (kg/m$^2$)*

*[2]Estimates obtained for the causal effect of BMI on mortality from malignant melanoma in males only used 76/77 SNPs, as all men who died from malignant melanoma had a dosage of '0' for one SNP (rs17024393); thus, providing no variation in mortality risk with this SNP.*

Table S8c. MR-Egger analysis of BMI on all-cause and cause-specific mortality in UK Biobank females of White British ancestry

| **Cause of death** | **IVW** | | **Intercept of MR-Egger** | | **MR-Egger** | | **Weighted median** | | **Weighted mode** | |
|---|---|---|---|---|---|---|---|---|---|---|
| | **HR (95% CI)[1]** | ***P*** | **HR (95% CI)[1]** | ***P*** | **HR (95% CI)[1]** | ***P*** | **HR (95% CI)[1]** | ***P*** | **HR (95% CI)[1]** | ***P*** |
| All-cause | 1.02 (0.96, 1.08) | 0.50 | 1.01 (0.99, 1.03) | 0.27 | 0.95 (0.82, 1.09) | 0.45 | 1.00 (0.93, 1.09) | 0.83 | 1.01 (0.92, 1.12) | 0.80 |
| Cardiovascular disease | 1.09 (0.94, 1.26) | 0.27 | 1.03 (0.98, 1.08) | 0.23 | 0.88 (0.61, 1.28) | 0.51 | 0.90 (0.80, 1.22) | 0.89 | 0.85 (0.64, 1.12) | 0.25 |
| Coronary heart disease | 1.10 (0.88, 1.37) | 0.40 | 1.03 (0.96, 1.11) | 0.34 | 0.86 (0.50, 1.49) | 0.58 | 1.05 (0.74, 1.45) | 0.82 | 1.25 (0.81, 1.93) | 0.31 |
| Stroke | 0.95 (0.72, 1.24) | 0.68 | 1.05 (0.96, 1.14) | 0.31 | 0.69 (0.35, 1.35) | 0.27 | 0.85 (0.61, 1.27) | 0.48 | 0.87 (0.54, 1.41) | 0.57 |
| Other cardiovascular diseases | 1.31 (1.01, 1.71) | 0.05 | 1.01 (0.93, 1.10) | 0.85 | 1.24 (0.64, 2.40) | 0.53 | 1.29 (0.86, 1.96) | 0.27 | 1.43 (0.84, 2.44) | 0.19 |
| Respiratory diseases | 1.03 (0.83, 1.28) | 0.80 | 1.03 (0.96, 1.11) | 0.39 | 0.83 (0.48, 1.43) | 0.49 | 0.87 (0.65, 1.35) | 0.68 | 0.89 (0.56, 1.39) | 0.6 |
| Cancer | 0.98 (0.92, 1.04) | 0.54 | 1.01 (0.99, 1.03) | 0.25 | 0.91 (0.78, 1.05) | 0.19 | 0.96 (0.89, 1.08) | 0.63 | 0.97 (0.87, 1.09) | 0.61 |
| Lung cancer | 1.02 (0.88, 1.17) | 0.83 | 1.02 (0.98, 1.07) | 0.34 | 0.87 (0.62, 1.23) | 0.43 | 0.95 (0.78, 1.19) | 0.70 | 0.90 (0.68, 1.21) | 0.51 |
| Breast cancer | 0.87 (0.76, 0.99) | 0.03 | 0.98 (0.94, 1.03) | 0.41 | 0.98 (0.71, 1.37) | 0.92 | 0.92 (0.73, 1.10) | 0.34 | 0.92 (0.70, 1.21) | 0.54 |
| Pre-menopausal | 0.83 (0.55, 1.27) | 0.39 | 0.99 (0.86, 1.13) | 0.85 | 0.92 (0.32, 2.61) | 0.87 | 0.94 (0.45, 1.65) | 0.69 | 1.01 (0.46, 2.22) | 0.98 |
| Post-menopausal | 0.87 (0.76, 1.00) | 0.05 | 0.98 (0.94, 1.03) | 0.42 | 0.99 (0.70, 1.41) | 0.97 | 0.91 (0.72, 1.13) | 0.35 | 0.92 (0.68, 1.23) | 0.57 |
| Colorectal cancer | 1.01 (0.80, 1.28) | 0.92 | 1.03 (0.95, 1.11) | 0.51 | 0.85 (0.47, 1.53) | 0.58 | 1.26 (0.81, 1.52) | 0.44 | 1.30 (0.82, 2.05) | 0.26 |
| Pancreatic cancer | 1.02 (0.82, 1.27) | 0.87 | 1.03 (0.96, 1.11) | 0.38 | 0.81 (0.47, 1.41) | 0.46 | 1.08 (0.73, 1.47) | 0.79 | 1.12 (0.76, 1.65) | 0.58 |
| Ovarian cancer | 1.16 (0.94, 1.42) | 0.16 | 1.03 (0.96, 1.10) | 0.39 | 0.95 (0.57, 1.57) | 0.83 | 1.17 (0.83, 1.55) | 0.45 | 1.20 (0.82, 1.76) | 0.35 |
| Endometrial cancer | 0.73 (0.49, 1.11) | 0.14 | 1.02 (0.90, 1.17) | 0.73 | 0.62 (0.22, 1.76) | 0.36 | 0.58 (0.33, 1.12) | 0.12 | 0.43 (0.17, 1.05) | 0.07 |
| Oesophageal cancer | 1.03 (0.70, 1.51) | 0.88 | 1.03 (0.91, 1.17) | 0.60 | 0.82 (0.31, 2.13) | 0.67 | 0.89 (0.48, 1.59) | 0.70 | 0.83 (0.35, 1.97) | 0.68 |
| Malignant melanoma | 1.44 (0.91, 2.26) | 0.11 | 1.07 (0.92, 1.24) | 0.37 | 0.89 (0.29, 2.78) | 0.85 | 1.41 (0.58, 2.14) | 0.75 | 1.06 (0.41, 2.71) | 0.91 |
| Kidney cancer | 0.77 (0.50, 1.18) | 0.22 | 1.04 (0.90, 1.19) | 0.58 | 0.58 (0.20, 1.70) | 0.32 | 0.84 (0.38, 1.58) | 0.57 | 0.80 (0.29, 2.21) | 0.67 |
| Brain cancer | 0.90 (0.66, 1.22) | 0.50 | 1.00 (0.90, 1.10) | 0.93 | 0.93 (0.43, 2.00) | 0.85 | 1.03 (0.62, 1.69) | 0.88 | 1.05 (0.58, 1.90) | 0.87 |
| Liver cancer | 0.96 (0.67, 1.39) | 0.84 | 1.10 (0.98, 1.24) | 0.10 | 0.47 (0.19, 1.18) | 0.11 | 0.67 (0.41, 1.25) | 0.27 | 0.47 (0.22, 1.01) | 0.06 |
| Lymphatic cancer | 1.05 (0.86, 1.29) | 0.61 | 1.02 (0.96, 1.09) | 0.53 | 0.91 (0.55, 1.51) | 0.70 | 0.91 (0.70, 1.30) | 0.79 | 0.86 (0.55, 1.33) | 0.49 |
| Other cancers | 1.03 (0.88, 1.22) | 0.69 | 0.99 (0.94, 1.04) | 0.70 | 1.11 (0.74, 1.68) | 0.61 | 1.07 (0.85, 1.41) | 0.44 | 1.10 (0.77, 1.57) | 0.60 |
| External causes | 1.59 (1.19, 2.12) | 0.002 | 1.01 (0.92, 1.11) | 0.77 | 1.44 (0.71, 2.94) | 0.31 | 1.40 (0.89, 2.26) | 0.10 | 1.37 (0.78, 2.42) | 0.28 |
| Other | 1.06 (0.92, 1.22) | 0.43 | 0.97 (0.93, 1.02) | 0.23 | 1.29 (0.91, 1.83) | 0.16 | 1.11 (0.94, 1.43) | 0.18 | 1.29 (0.95, 1.77) | 0.11 |

*BMI = body mass index; CI = confidence interval; HR = hazard ratio; IVW = inverse variance weighted; MR = Mendelian randomization*

*[1]Adjusted for secular trends (date of birth) and the first ten genetic principal components, estimates represent HR with each unit increase in BMI ($kg/m^2$)*

Table S9. MR analyses of all-cause and cause-specific mortality by BMI in UK Biobank participants of White British ancestry additionally adjusting for UK BiLEVE[1]

| **Cause of death** | **Whole sample** | | **Males** | | **Females** | |
|---|---|---|---|---|---|---|
| | **HR (95% CI)[2]** | ***P*-value** | **HR (95% CI)[2]** | ***P*-value** | **HR (95% CI)[2]** | ***P*-value** |
| All-cause | 1.02 (0.97, 1.06) | 0.46 | 1.02 (0.97, 1.09) | 0.41 | 1.00 (0.94, 1.07) | 0.91 |
| Cardiovascular disease | 1.12 (1.02, 1.23) | 0.02 | 1.10 (0.98, 1.24) | 0.10 | 1.18 (0.98, 1.43) | 0.09 |
| Coronary heart disease | 1.19 (1.04, 1.35) | 0.01 | 1.20 (1.04, 1.40) | 0.02 | 1.19 (0.86, 1.67) | 0.29 |
| Stroke | 0.98 (0.78, 1.24) | 0.88 | 0.90 (0.64, 1.26) | 0.52 | 1.10 (0.78, 1.54) | 0.59 |
| Aortic aneurysm | 0.86 (0.59, 1.27) | 0.46 | 0.82 (0.52, 1.31) | 0.41 | - | - |
| Other cardiovascular disease | 1.13 (0.93, 1.38) | 0.22 | 1.06 (0.82, 1.37) | 0.66 | 1.30 (0.93, 1.83) | 0.13 |
| Respiratory diseases | 0.88 (0.72, 1.07) | 0.20 | 0.84 (0.64, 1.09) | 0.18 | 0.96 (0.68, 1.34) | 0.79 |
| Cancer | 0.98 (0.93, 1.03) | 0.41 | 1.00 (0.92, 1.08) | 0.94 | 0.96 (0.89, 1.03) | 0.25 |
| Lung cancer | 0.96 (0.84, 1.11) | 0.60 | 0.96 (0.78, 1.17) | 0.68 | 0.97 (0.79, 1.19) | 0.76 |
| Prostate cancer | - | - | 0.73 (0.57, 0.93) | 0.01 | - | - |
| Breast cancer | - | - | - | - | 0.87 (0.73, 1.02) | 0.09 |
| Pre-menopausal | - | - | - | - | 0.81 (0.50, 1.30) | 0.38 |
| Post-menopausal | - | - | - | - | 0.87 (0.73, 1.04) | 0.14 |
| Colorectal cancer | 1.10 (0.93, 1.29) | 0.25 | 1.17 (0.93, 1.47) | 0.18 | 1.02 (0.81, 1.30) | 0.84 |
| Pancreatic cancer | 1.02 (0.83, 1.25) | 0.84 | 1.04 (0.77, 1.40) | 0.81 | 1.01 (0.76, 1.33) | 0.97 |
| Stomach cancer | 1.30 (0.91, 1.87) | 0.16 | 1.32 (0.84, 2.08) | 0.23 | - | - |
| Ovarian cancer | - | - | - | - | 1.13 (0.88, 1.46) | 0.33 |
| Endometrial cancer | - | - | - | - | 0.71 (0.41, 1.23) | 0.23 |
| Oesophageal cancer | 1.08 (0.84, 1.38) | 0.55 | 1.20 (0.88, 1.62) | 0.25 | 0.81 (0.50, 1.31) | 0.39 |
| Malignant melanoma | 1.06 (0.76, 1.50) | 0.72 | 0.91 (0.57, 1.46) | 0.71 | 1.33 (0.78, 2.27) | 0.29 |
| Kidney cancer | 0.96 (0.70, 1.32) | 0.81 | 1.08 (0.73, 1.59) | 0.70 | 0.71 (0.39, 1.30) | 0.27 |
| Bladder cancer | 0.84 (0.56, 1.26) | 0.40 | 0.90 (0.55, 1.46) | 0.66 | - | - |
| Brain cancer | 1.01 (0.80, 1.27) | 0.95 | 1.30 (0.93, 1.80) | 0.12 | 0.75 (0.53, 1.04) | 0.09 |
| Liver cancer | 0.95 (0.69, 1.30) | 0.74 | 0.81 (0.52, 1.25) | 0.34 | 1.17 (0.73, 1.86) | 0.51 |
| Lymphatic cancer | 1.03 (0.86, 1.22) | 0.76 | 1.04 (0.82, 1.32) | 0.74 | 1.01 (0.77, 1.32) | 0.94 |
| Other cancer | 0.95 (0.82, 1.10) | 0.52 | 0.90 (0.74, 1.11) | 0.32 | 1.02 (0.82, 1.27) | 0.87 |
| External causes | 1.35 (1.09, 1.67) | 0.01 | 1.25 (0.94, 1.66) | 0.13 | 1.56 (1.10, 2.21) | 0.01 |
| Other | 1.00 (0.88, 1.14) | 0.97 | 1.00 (0.84, 1.18) | 0.97 | 1.01 (0.83, 1.23) | 0.92 |

*BMI = body mass index; CI = confidence interval; HR = hazard ratio*

*[1]There was evidence of differential array effect on markers scattered across the genome; therefore, the UK BiLEVE study genotyped on the Affymetrix Axiom Array was considered as a confounder*

*[2]Adjusted for secular trends (date of birth), highest household occupation, education, smoking status, alcohol intake, physical activity, the first ten genetic principal components and genotyping chip, estimates represent HR with each unit increase in BMI (kg/m$^2$)*

Table S10. Association between weighted GRS (comprising 70 SNPs) and BMI in UK Biobank participants of White British ancestry

| **Sample** | **N** | **Effect estimate (95% CI)**[1] | ***P*-value** | **$R^2$ (%)**[2] |
|---|---|---|---|---|
| Whole sample | 335,308 | 0.11 (0.11, 0.11) | $<1.20 \times 10^{-307}$ | 1.67 |
| Males | 154,967 | 0.10 (0.10, 0.11) | $<1.20 \times 10^{-307}$ | 1.90 |
| Females | 180,341 | 0.12 (0.11, 0.12) | $<1.20 \times 10^{-307}$ | 1.57 |

*BMI = body mass index; CI = confidence interval; GRS = genetic risk score; SNP = single nucleotide polymorphism*

[1]*Effect estimate (and corresponding P-value) represents the change in BMI (kg/m$^2$) per BMI-increasing allele in individuals of White British ancestry adjusted for the first ten genetic principal components*

[2]*Variance explained*

Table S11. MR analyses of all-cause and cause-specific mortality by BMI in UK Biobank participants of White British ancestry using a GRS restricted to variants known to be classified as having a secondary signal within a locus to other phenotypes

| **Cause of death** | **Whole sample** | | **Males** | | **Females** | |
|---|---|---|---|---|---|---|
| | **HR (95% CI)[2]** | ***P*-value** | **HR (95% CI)[2]** | ***P*-value** | **HR (95% CI)[2]** | ***P*-value** |
| All-cause | 1.01 (0.97, 1.06) | 0.60 | 1.02 (0.96, 1.08) | 0.53 | 1.00 (0.94, 1.07) | 0.95 |
| Cardiovascular disease | 1.12 (1.02, 1.24) | 0.02 | 1.10 (0.98, 1.25) | 0.11 | 1.20 (0.99, 1.46) | 0.06 |
| Coronary heart disease | 1.21 (1.06, 1.39) | 0.01 | 1.21 (1.04, 1.42) | 0.02 | 1.31 (0.93, 1.85) | 0.12 |
| Stroke | 0.97 (0.76, 1.24) | 0.84 | 0.94 (0.66, 1.33) | 0.73 | 1.02 (0.72, 1.44) | 0.92 |
| Aortic aneurysm | 0.89 (0.60, 1.33) | 0.58 | 0.87 (0.54, 1.41) | 0.57 | - | - |
| Other cardiovascular disease | 1.10 (0.90, 1.36) | 0.34 | 0.99 (0.76, 1.29) | 0.95 | 1.38 (0.97, 1.95) | 0.07 |
| Respiratory diseases | 0.88 (0.72, 1.09) | 0.24 | 0.83 (0.64, 1.09) | 0.19 | 0.97 (0.69, 1.38) | 0.88 |
| Cancer | 0.97 (0.91, 1.02) | 0.22 | 0.98 (0.91, 1.07) | 0.70 | 0.95 (0.88, 1.02) | 0.16 |
| Lung cancer | 0.96 (0.83, 1.11) | 0.62 | 0.97 (0.78, 1.19) | 0.75 | 0.96 (0.78, 1.19) | 0.73 |
| Prostate cancer | - | - | 0.68 (0.53, 0.88) | 0.004 | - | - |
| Breast cancer | - | - | - | - | 0.86 (0.72, 1.02) | 0.08 |
| Pre-menopausal | - | - | - | - | 0.88 (0.54, 1.44) | 0.61 |
| Post-menopausal | - | - | - | - | 0.85 (0.71, 1.03) | 0.09 |
| Colorectal cancer | 1.07 (0.90, 1.26) | 0.44 | 1.14 (0.90, 1.44) | 0.29 | 1.00 (0.78, 1.27) | 0.98 |
| Pancreatic cancer | 0.96 (0.78, 1.19) | 0.73 | 1.03 (0.75, 1.40) | 0.87 | 0.91 (0.68, 1.21) | 0.50 |
| Stomach cancer | 1.31 (0.90, 1.90) | 0.16 | 1.34 (0.84, 2.14) | 0.22 | - | - |
| Ovarian cancer | - | - | - | - | 1.13 (0.87, 1.46) | 0.36 |
| Endometrial cancer | - | - | - | - | 0.76 (0.43, 1.34) | 0.34 |
| Oesophageal cancer | 1.13 (0.87, 1.46) | 0.35 | 1.26 (0.92, 1.72) | 0.15 | 0.84 (0.51, 1.38) | 0.49 |
| Malignant melanoma | 1.10 (0.77, 1.57) | 0.60 | 0.94 (0.58, 1.52) | 0.80 | 1.39 (0.80, 2.42) | 0.24 |
| Kidney cancer | 0.97 (0.70, 1.34) | 0.84 | 1.04 (0.69, 1.56) | 0.85 | 0.81 (0.43, 1.50) | 0.49 |
| Bladder cancer | 0.82 (0.54, 1.25) | 0.35 | 0.87 (0.53, 1.43) | 0.58 | - | - |
| Brain cancer | 1.03 (0.81, 1.31) | 0.81 | 1.31 (0.93, 1.84) | 0.12 | 0.78 (0.55, 1.10) | 0.15 |
| Liver cancer | 0.90 (0.65, 1.24) | 0.52 | 0.80 (0.51, 1.26) | 0.33 | 1.05 (0.65, 1.69) | 0.85 |
| Lymphatic cancer | 1.00 (0.84, 1.20) | 0.99 | 1.06 (0.83, 1.36) | 0.64 | 0.92 (0.70, 1.22) | 0.56 |
| Other cancer | 0.95 (0.81, 1.11) | 0.50 | 0.87 (0.70, 1.07) | 0.18 | 1.07 (0.85, 1.34) | 0.58 |
| External causes | 1.35 (1.08, 1.69) | 0.01 | 1.28 (0.95, 1.72) | 0.11 | 1.53 (1.06, 2.19) | 0.02 |
| Other | 1.02 (0.90, 1.16) | 0.75 | 1.00 (0.84, 1.19) | 0.99 | 1.05 (0.86, 1.29) | 0.62 |

*BMI = body mass index; CI = confidence interval; HR = hazard ratio*

*[1]There was evidence of differential array effect on markers scattered across the genome; therefore, the UK BiLEVE study genotyped on the Affymetrix Axiom Array was considered as a confounder*

*[2]Adjusted for secular trends (date of birth), highest household occupation, education, smoking status, alcohol intake, physical activity, the first ten genetic principal components and genotyping chip, estimates represent HR with each unit increase in BMI (kg/m$^2$)*

Figure S1. MR-Egger analysis for mortality outcomes with any evidence for pleiotropy

(a) other cancers in the whole UK Biobank sample

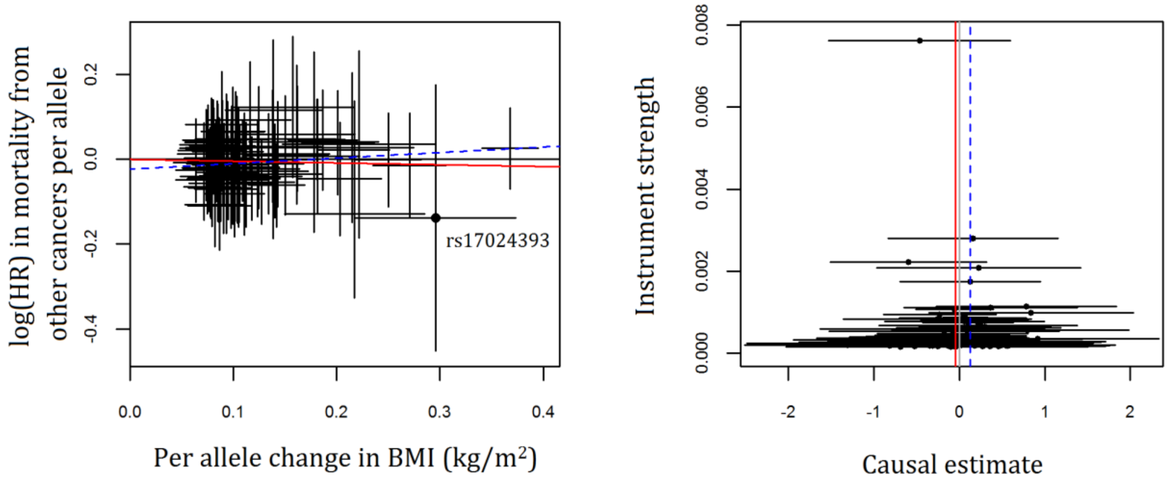


(b) other causes in the whole UK Biobank sample

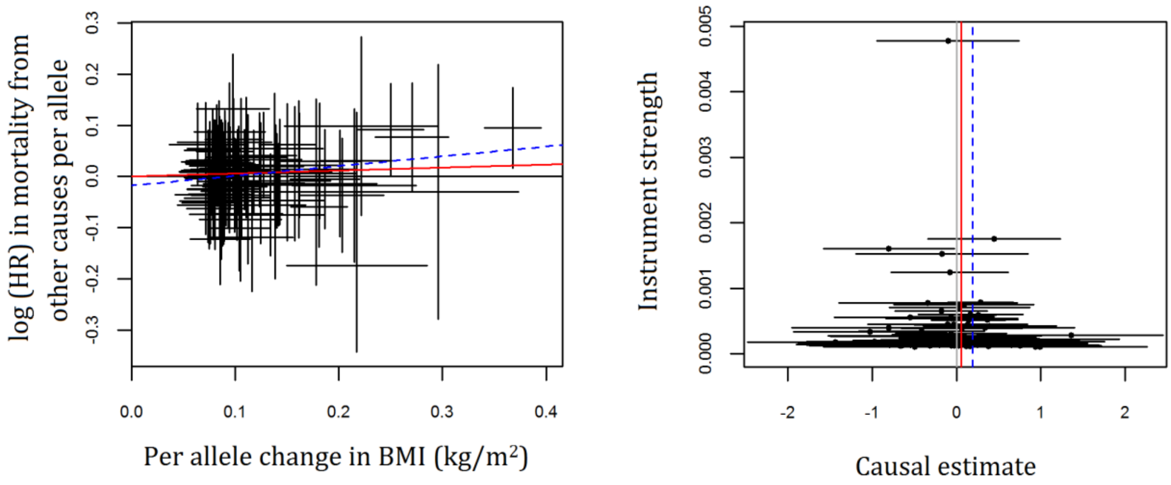

Figure S2. MR-Egger analysis for mortality outcomes with any evidence for pleiotropy in sex-stratified analyses

(a) other cancers in males only

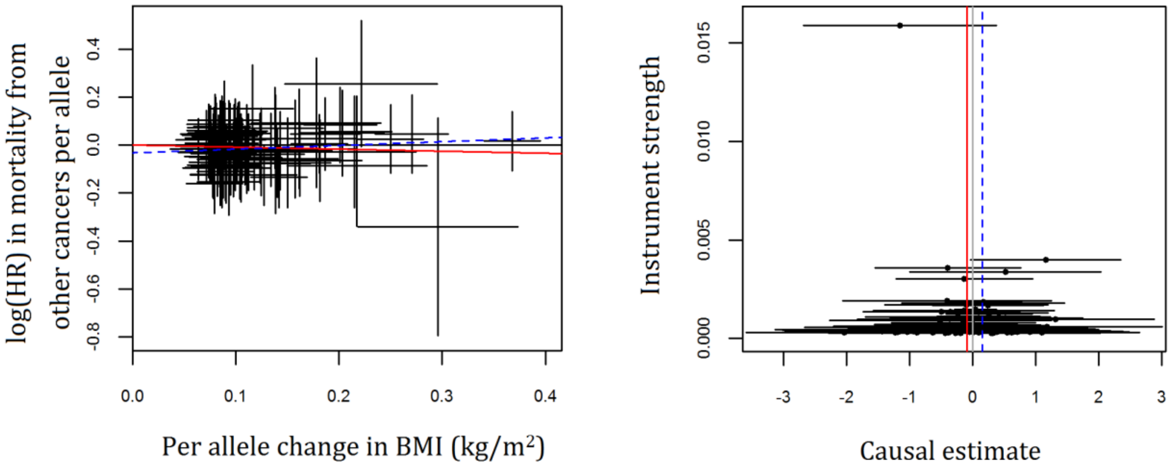


(b) other causes in males only

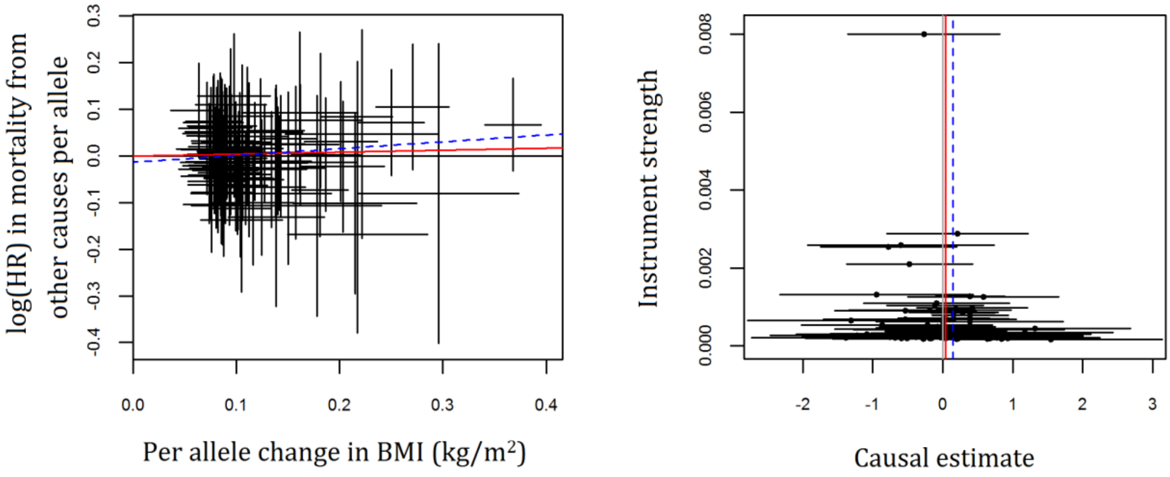

2. Locke AE, Kahali B, Berndt SI, Justice AE, Pers TH, Day FR, Powell C, Vedantam S, Buchkovich ML, Yang J, Croteau-Chonka DC, Esko T, Fall T, Ferreira T, Gustafsson S, Kutalik Z, Luan Ja, Mägi R, Randall JC, Winkler TW, Wood AR, Workalemahu T, Faul JD, Smith JA, Zhao JH, Zhao W, Chen J, Fehrmann R, Hedman ÅK, Karjalainen J, Schmidt EM, Absher D, Amin N, Anderson D, Beekman M, Bolton JL, Bragg-Gresham JL, Buyske S, Demirkan A, Deng G, Ehret GB, Feenstra B, Feitosa MF, Fischer K, Goel A, Gong J, Jackson AU, Kanoni S, Kleber ME, Kristiansson K, Lim U, Lotay V, Mangino M, Leach IM, Medina-Gomez C, Medland SE, Nalls MA, Palmer CD, Pasko D, Pechlivanis S, Peters MJ, Prokopenko I, Shungin D, Stančáková A, Strawbridge RJ, Sung YJ, Tanaka T, Teumer A, Trompet S, van der Laan SW, van Setten J, Van Vliet-Ostaptchouk JV, Wang Z, Yengo L, Zhang W, Isaacs A, Albrecht E, Ärnlöv J, Arscott GM, Attwood AP, Bandinelli S, Barrett A, Bas IN, Bellis C, Bennett AJ, Berne C, Blagieva R, Blüher M, Böhringer S, Bonnycastle LL, Böttcher Y, Boyd HA, Bruinenberg M, Caspersen IH, Chen Y-DI, Clarke R, Daw EW, de Craen AJM, Delgado G, Dimitriou M, Doney ASF, Eklund N, Estrada K, Eury E, Folkersen L, Fraser RM, Garcia ME, Geller F, Giedraitis V, Gigante B, Go AS, Golay A, Goodall AH, Gordon SD, Gorski M, Grabe H-J, Grallert H, Grammer TB, Gräßler J, Grönberg H, Groves CJ, Gusto G, Haessler J, Hall P, Haller T, Hallmans G, Hartman CA, Hassinen M, Hayward C, Heard-Costa NL, Helmer Q, Hengstenberg C, Holmen O, Hottenga J-J, James AL, Jeff JM, Johansson Å, Jolley J, Juliusdottir T, Kinnunen L, Koenig W, Koskenvuo M, Kratzer W, Laitinen J, Lamina C, Leander K, Lee NR, Lichtner P, Lind L, Lindström J, Lo KS, Lobbens S, Lorbeer R, Lu Y, Mach F, Magnusson PKE, Mahajan A, McArdle WL, McLachlan S, Menni C, Merger S, Mihailov E, Milani L, Moayyeri A, Monda KL, Morken MA, Mulas A, Müller G, Müller-Nurasyid M, Musk AW, Nagaraja R, Nöthen MM, Nolte IM, Pilz S, Rayner NW, Renstrom F, Rettig R, Ried JS, Ripke S, Robertson NR, Rose LM, Sanna S, Scharnagl H, Scholtens S, Schumacher FR, Scott WR, Seufferlein T, Shi J, Smith AV, Smolonska J, Stanton AV, Steinthorsdottir V, Stirrups K, Stringham HM, Sundström J, Swertz MA, Swift AJ, Syvänen A-C, Tan S-T, Tayo BO, Thorand B, Thorleifsson G, Tyrer JP, Uh H-W, Vandenput L, Verhulst FC, Vermeulen SH, Verweij N, Vonk JM, Waite LL, Warren HR, Waterworth D, Weedon MN, Wilkens LR, Willenborg C, Wilsgaard T, Wojczynski MK, Wong A, Wright AF, Zhang Q, The LifeLines Cohort S, Brennan EP, Choi M, Dastani Z, Drong AW, Eriksson P, Franco-Cereceda A, Gådin JR, Gharavi AG, Goddard ME, Handsaker RE, Huang J, Karpe F, Kathiresan S, Keildson S, Kiryluk K, Kubo M, Lee J-Y, Liang L, Lifton RP, Ma B, McCarroll SA, McKnight AJ, Min JL, Moffatt MF, Montgomery GW, Murabito JM, Nicholson G, Nyholt DR, Okada Y, Perry JRB, Dorajoo R, Reinmaa E, Salem RM, Sandholm N, Scott RA, Stolk L, Takahashi A, Tanaka T, van 't Hooft FM, Vinkhuyzen AAE, Westra H-J, Zheng W, Zondervan KT, The AC, The A-BMIWG, The CDC, The CC, The G, The I, The MI, The Mu TC, The MC, The PC, The ReproGen C, The GC, The International Endogene C, Heath AC, Arveiler D, Bakker SJL, Beilby J, Bergman RN, Blangero J, Bovet P, Campbell H, Caulfield MJ, Cesana G, Chakravarti A, Chasman DI, Chines PS, Collins FS, Crawford DC, Cupples LA, Cusi D, Danesh J, de Faire U, den Ruijter HM, Dominiczak AF, Erbel R, Erdmann J, Eriksson JG, Farrall M, Felix SB, Ferrannini E, Ferrières J, Ford I, Forouhi NG, Forrester T, Franco OH, Gansevoort RT, Gejman PV, Gieger C, Gottesman O, Gudnason V, Gyllensten U, Hall AS, Harris TB, Hattersley AT, Hicks AA, Hindorff LA, Hingorani AD, Hofman A, Homuth G, Hovingh GK, Humphries SE, Hunt SC, Hyppönen E, Illig T, Jacobs KB, Jarvelin M-R, Jöckel K-H, Johansen B, Jousilahti P, Jukema JW, Jula AM, Kaprio J, Kastelein JJP, Keinanen-Kiukaanniemi SM, Kiemeney LA, Knekt P, Kooner JS, Kooperberg C, Kovacs P, Kraja AT, Kumari M, Kuusisto J, Lakka TA, Langenberg C, Marchand LL, Lehtimäki T, Lyssenko V, Männistö S, Marette A, Matise TC, McKenzie CA, McKnight B, Moll FL, Morris AD, Morris AP, Murray JC, Nelis M, Ohlsson C, Oldehinkel AJ, Ong KK, Madden PAF, Pasterkamp G, Peden JF, Peters A, Postma DS, Pramstaller PP, Price JF, Qi L, Raitakari OT, Rankinen T, Rao DC, Rice TK, Ridker PM, Rioux JD, Ritchie MD, Rudan I, Salomaa V, Samani NJ, Saramies J, Sarzynski MA, Schunkert H, Schwarz PEH, Sever P, Shuldiner AR, Sinisalo J, Stolk RP, Strauch K, Tönjes A, Trégouët D-A, Tremblay A, Tremoli E, Virtamo J, Vohl M-C, Völker U, Waeber G, Willemsen G, Witteman JC, Zillikens MC, Adair LS, Amouyel P, Asselbergs FW, Assimes TL, Bochud M, Boehm BO, Boerwinkle E, Bornstein SR, Bottinger EP, Bouchard C, Cauchi S, Chambers JC, Chanock SJ, Cooper RS, de Bakker PIW, Dedoussis G, Ferrucci L, Franks PW, Froguel P, Groop LC, Haiman CA, Hamsten A, Hui J, Hunter DJ, Hveem K, Kaplan RC, Kivimaki M, Kuh D, Laakso M, Liu Y, Martin NG, März W, Melbye M, Metspalu A, Moebus S, Munroe PB, Njølstad I, Oostra BA, Palmer CNA, Pedersen NL, Perola M, Pérusse L, Peters U, Power C, Quertermous T, Rauramaa R, Rivadeneira F, Saaristo TE, Saleheen D, Sattar N, Schadt EE, Schlessinger D, Slagboom PE, Snieder H, Spector TD, Thorsteinsdottir U, Stumvoll M, Tuomilehto J, Uitterlinden AG, Uusitupa M, van der Harst P, Walker M, Wallaschofski H, Wareham NJ, Watkins H, Weir DR, Wichmann HE, Wilson JF, Zanen P, Borecki IB, Deloukas P, Fox CS, Heid IM, O'Connell JR, Strachan DP, Stefansson K, van Duijn CM, Abecasis GR, Franke L, Frayling TM, McCarthy MI, Visscher PM, Scherag A, Willer CJ, Boehnke M, Mohlke KL,

Lindgren CM, Beckmann JS, Barroso I, North KE, Ingelsson E, Hirschhorn JN, Loos RJF and Speliotes EK. Genetic studies of body mass index yield new insights for obesity biology. *Nature*. 2015;518:197-206.
3. Corbin LJ, Richmond RC, Wade KH, Burgess S, Bowden J, Smith GD and Timpson NJ. BMI as a Modifiable Risk Factor for Type 2 Diabetes: Refining and Understanding Causal Estimates Using Mendelian Randomization. *Diabetes*. 2016;65:3002-3007.
4. Tyrrell J, Jones SE, Beaumont R, Astley CM, Lovell R, Yaghootkar H, Tuke M, Ruth KS, Freathy RM, Hirschhorn JN, Wood AR, Murray A, Weedon MN and Frayling TM. Height, body mass index, and socioeconomic status: mendelian randomisation study in UK Biobank. *BMJ*. 2016;352.
5. Yaghootkar H, Lotta LA, Tyrrell J, Smit RAJ, Jones SE, Donnelly L, Beaumont R, Campbell A, Tuke MA, Hayward C, Ruth KS, Padmanabhan S, Jukema JW, Palmer CC, Hattersley A, Freathy RM, Langenberg C, Wareham NJ, Wood AR, Murray A, Weedon MN, Sattar N, Pearson E, Scott RA and Frayling TM. Genetic Evidence for a Link Between Favorable Adiposity and Lower Risk of Type 2 Diabetes, Hypertension, and Heart Disease. *Diabetes*. 2016;65:2448.
